# Derivation of cardiomyocyte-propelled motile aggregates from stem cells

**DOI:** 10.1101/2025.07.09.663178

**Authors:** Christine Ho, Fokion Glykofrydis, Gaveen Godage, Kyle Poon, Minnal Kunnan, Benjamin Swedlund, Sandra Murillo, Leonardo Morsut

## Abstract

Robotics draws inspiration from biology, particularly animal locomotion based on muscle-driven contractions. While traditional engineering assembles components sequentially, locomotive animals are built via self-organized developmental programs. Stem cells, under the right conditions, can mimic these processes in vitro, offering a pathway to develop muscle-propelled biobots in a self-organized building process. Here, we demonstrate that existent cardiogenic gastruloid protocols can produce motile aggregates from mouse embryonic stem cells, although with very limited efficiency. We then identify a novel protocol that yields contractile aggregates with higher frequency and larger contractile areas. In this novel protocol, mesendoderm induction using TGF-beta ligands is followed by cardiogenic induction with FGFs and VEGF. Synthetic organizers further control contraction localization. Aggregates developed via this protocol show enhanced motility, marking a step forward towards building motile cardiobots from self-organized biological material. This strategy opens new possibilities for designing autonomous biobots and studying the evolution of muscle-powered movement of multicellular organisms and cardiovascular development.

## Introduction

To what extent can we control multicellular systems development toward user-defined shapes and functions? This question lies at the intersection of key challenges in evolutionary, developmental, cell, and synthetic biology, as well as tissue engineering and regenerative medicine. This emerging multidisciplinary endeavor to steer the collective behavior of cells has captured significant interest for two primary reasons. First, it presents a powerful engineering paradigm for achieving complex biological outcomes offering transformative potential for creating sophisticated tissues in regenerative medicine and other biomedical applications. Second, by using a constructive approach, orthogonal to the classic analytical approach of biology, it provides a novel angle towards the understanding of the design rules of both synthetic and natural multicellular systems. Over the past decade, there has been a sharp rise in interest around the construction of biological structures. Among these initiatives, a subset of functional, bioengineered assemblies has led to the emergence of a distinct class of motile synthetic systems known as biobots.

### Building biobots with scaffold

Early iterations of biobots were generated combining synthetic scaffolds with cell-based actuators, such as muscles, nerves, and neuromuscular junctions, as well as engineered cell lines with programmable properties^1–3^. With this approach, microswimmers, walkers and fluidic devices could be built, targeting applications in drug delivery and therapeutics, and innovations in microfluidics, manufacturing, bio-electronic interfaces, control and communication^4–9^. In these cases, the living elements are embedded within precisely designed 3D architectures to enhance and leverage their inherent biological activity. One limitation of this approach is that the fabrication approach is top-down and requires external intervention, leading to a limited number of prototypes that can be tested and iteratively improved, thus limiting the exploration of different designs and comparison of performance.

### The evolutionary case of animal embryonic development

In nature, multicellular systems that display high-level emergent collective functions, in particular motility, are built with a different paradigm: through the process of embryonic development, the process by which a single cell (e.g. fertilized oocyte) gives rise to a multicellular organism. This continuous process is self-organized and does not require external scaffolds. In animals, the resulting organism has several features like patterning, motility, symmetry, information processing, energy acquisition and utilization, that allow them to efficiently interact with their environment. These features emerged during evolution from precursors that were unicellular; the evolution of the development of motility in multicellular structures has been a center of several intriguing studies and theories ^10–20^. It is widely believed that an initial cilia-based process was substituted by a muscle-contraction based process, likely evolving from more ancestral cytoskeletal contraction and cell division machinery ^21–23^. All contractile cells in modern-day animals, e.g. smooth muscle, skeletal muscle and cardiac muscle, are postulated to have evolved from this ancestral mechanism that supported early muscle-driven motility. The architecture of the ancestral animal is unknown, but from a survey of extant ancestral animals, it most likely contained a high percentage of muscle tissue^23^ (**Fig. 1A**). Whether we can, using modern-day cells, derive aggregates that undergo development and become motile via muscle-propulsion has not yet been explored. The goal of this study is to define a direct developmental trajectory that takes stem/progenitor cells into a multi-lineage differentiation to generate aggregates with autonomous contractility that generate enough force to displace the aggregate and make them motile (**Fig.1B**).

**Figure 1.**
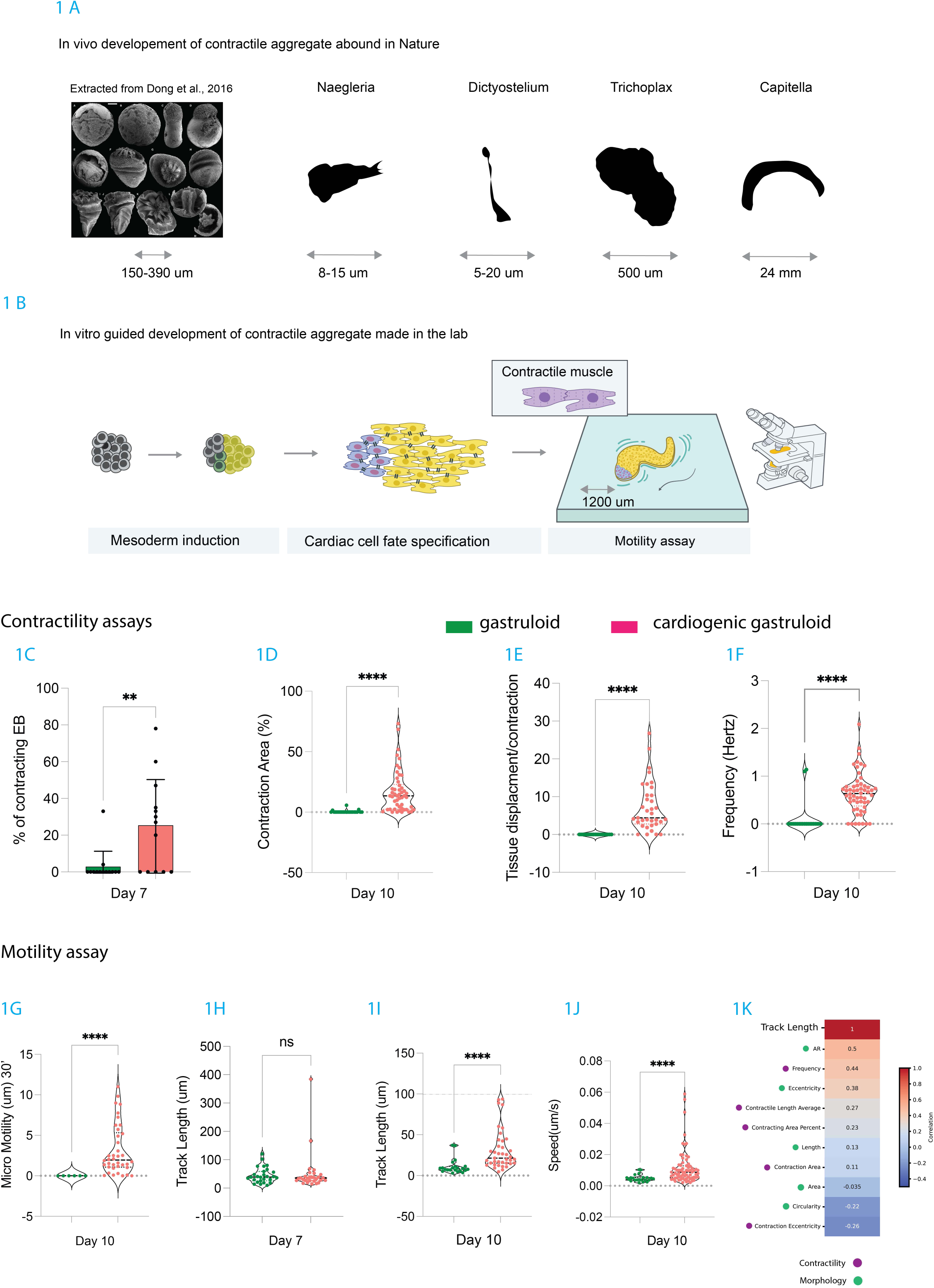
Existing cardiogenic gastruloid protocols generate aggregates that exhibit motility only in contractile samples in low efficiency. **A.** Examples of natural life forms that resemble contractile aggregates. First picture has been extracted from. ^70^. Relative scale bar represents: A, 151 μm; B, 149 μm; C, 204 μm; D, 183 μm; E, 217 μm; F, 208 μm; G, 212 μm; H, 225 μm; I, 388 μm; J, 348 μm; K, 257 μm; L, 192 μm; M, 211 μm.**B.** Schematic illustration of the guided development of contractile aggregates grown *in vitro*, adapted from the protocol described by Rossi et al., 2018 for cardiogenic gastruloids. Mouse embryonic stem cells (mESCs) are initially in a stem state (gray), and then differentiate towards mesoderm (dark yellow), and then to cardiomyocytes (light yellows). Other cell lineage are also present (represented in purple). A motility assay setup is shown on the far right, where multiple aggregates are placed together on a microscope surface. Starting from when contractions occur (depicted as after day 7 in the schematic) motility can be assayed. **C.** Bar graph showing the percentage of contracting embryoid bodies on Day 7 for both gastruloids (n=13) and cardiogenic gastruloids (n=13). *n* = number of individual aggregates. Statistical analysis was performed using the Mann–Whitney test. **D.** Violin plot showing the percentage of the total area of the aggregate that displays contractile features on Day 10 for gastruloids (n=22) and cardiogenic gastruloids (n=50). Dotted black line represents the median *n* = number of individual aggregates. Statistical analysis was performed using the Mann–Whitney test. **E.** Violin plot showing tissue displacement per contraction (in micrometers) on Day 10 in gastruloids (n=19) and cardiogenic gastruloids (n=35). See methods for quantification pipeline. *n* = number of individual aggregates. Dotted black line represents the median. Statistical analysis was performed using the Mann–Whitney test. **F.** Bar graph showing contraction frequency (in Hertz) on Day 10 for gastruloids (n=21) and cardiogenic gastruloids (n=56). Dotted black line represents the median. *n* = number of individual aggregates. Statistical analysis was performed using the Mann–Whitney test. **G.** Bar graph showing micro-motility (in micrometers) measured over 30 seconds in day 10 gastruloids (n=6) and cardiogenic gastruloids (n=38). Dotted black line represents the median *n* = number of individual aggregates. Statistical analysis was performed using the Mann–Whitney test. **H–I.** Bar graphs showing track length (in micrometers) during motility assays performed on Day 7 (H) and Day 10 (I) in gastruloids (n=22) and cardiogenic gastruloids (n=39). Dotted black line represents the median. *n* = number of individual aggregates. Statistical analysis was performed using the Mann–Whitney test. **J.** Bar graph showing speed (in micrometers/second) of individual embryoid bodies during motility assays on Day 10 for gastruloids (n=22) and cardiogenic gastruloids (n=72). Dotted black line represents the median. *n* = number of individual aggregates. Statistical analysis was performed using the Mann–Whitney test. **K.** Correlation map for cardiogenic gastruloids (n=39) showing the relationship between measured track length and the indicated morphological (green dot) and contractile (purple dot) features. The strength of correlation is indicated by shade of color, as shown in the correlation legend on the right. Statistical significance: *p < 0.05; **p < 0.01; ***p < 0.001.

### Building biobots without scaffold via embryonic development in vitro

Recent works have laid the foundation for attempting this goal. Motivated by regenerative medicine and developmental studies, the guided self-organization of stem cells into analogues of specific organs and tissues in vitro has seen successful progress in the last few decades ^24,25^. They can be targeted to specific cell types, or to fetal versions of organs known as organoids. In general, the protocols for deriving multicellular constructs from stem cells involve growth of stem cells in 2D or 3D and exposure to growth factor stimulation of morphogenetic signaling pathways. In early iterations of these protocols, growth factors came dissolved in the media; more recently, they started to come from microfluidic systems^26^, or from synthetic organizers ^27–29^. In the field of biobots, where the goal is not to recapitulate existing structures, but building new ones, first examples were motile Xenobots^30,31^ made of X.laevis cells, and anthrobots^32^, made with human stem cells. Both these biobots are motile, but move thanks to cilia-driven mechanisms. So far, none of the biobots developed with a developmental protocol move thanks to muscle-based contractions. Protocols to derive muscle cells in organoids context have been developed. A notable example are cardiogenic gastruloids which follow a developmental trajectory (i.e. continuous development from stem cells), and generate contractile cardiomyocytes ^33–35^. In these protocols, a brief 24-hour pulse of Wnt signaling induces mesoderm and gastrulation-like movements; if this is followed by cardiogenic stimulation with VEGF and FGF2 in N2B27 media, these structures continue their development towards cardiogenic mesoderm and finally cardiomyocytes-driven contractions can be seen ^36–39^. Whether the contractions can support motility has not been explored.

Here, we show that current cardiogenic gastruloid protocols produce motile aggregates, albeit at low efficiency and frequency. We study their behaviors and identify that limited contractile area is a limiting factor in their motility. Via a screening, we identified a protocol that generates contractile aggregates at higher frequency and with larger contractile areas, which we introduce as ‘cardiobots’. This protocol proceeds via a mesendoderm induction via Tgf-beta family ligands, followed by cardiogenic induction with FGFs and VEGF. We show how we can control shape and localization of contraction in these aggregates with synthetic organizers, where the morphogens are not dispersed in the media but provided by clusters of engineered cells, thus showing that the body plans of cardiobots can be controlled in space and time. Finally, we show that the soluble-factor based protocols generate aggregates with higher motility capacity. Two main morphotypes are identified, either elongated or a spherical shape, that are able to generate similar levels of motility. We characterize the features that support highest motility in cardiobots and identify contraction area, circularity, and contraction efficiency as key parameters for obtaining motile cardiobots. Collectively, our results show that cardiogenesis in a self-organization protocol can be improved with activin/BMP factors followed by FGFs/VEGF, and that this leads to aggregates that are capable of millimeter-scale motility. These protocols could therefore serve as a basis for derivation of efficient motile cardiobots for exploration, scavenging and regenerative applications, and also for the study of the evolution of the development of muscle-based multicellular motility and of the cardiovascular system.

## Results

### Cardiogenic gastruloid protocols generate motile aggregates at very low efficiency

We assessed if currently available cardiogenic gastruloid protocols generate aggregates capable of motility. To do so, we subjected mouse embryonic stem cells to gastruloid and cardiogenic gastruloid protocols ^35,40^. Briefly, 300 mESCs were seeded into U-bottom 96-well plates and cultured in N2B27 for 2 days. During this time, the cells aggregated to form embryoid bodies, which were then exposed to a 24-hour pulse of Chiron (Wnt signaling activator) to induce mesodermal differentiation. For conventional gastruloids, Chiron was washed out after the pulse, and the aggregates were cultured for an additional 3 days in fresh neurobasal supplement with B-27 and 2-2 supplements (N2B27) medium. For cardiogenic gastruloids, following Chiron removal on day 3, the aggregates were cultured for another 2 days in N2B27 supplemented with cardiogenic factors, including Ascorbic Acid (AA), Vascular Endothelial Growth Factor (VEGF), and Fibroblast Growth Factor 2 (FGF2) from day 4 to 6 post seeding. After 7 days of differentiation, the standard and cardiogenic gastruloids were transferred into low adherent 6-well plates on an orbital shaker from day 7 to day 10 (**Fig. S1A**). As expected, the use of cardiogenic medium between 96h (day 4) and 120h (day 5) led to a significant increase in the number of beating aggregates on day 7 of differentiation, compared to the standard gastruloid protocol. We observed an increase in the proportion of beating aggregates, ranging from 30% to 80% for different batches of differentiation (N=8, **Fig. 1C**), in line with what was reported previously^40^. To characterize the contractions of these aggregates, we measured their frequency, the area of contracting tissue, and the tissue displacement per contraction. To extract frequency and area of contracting tissue, we monitored the calcium activity spatially in the aggregates using Fluo-4, a cell permeant to label intracellular calcium, and live imaging (See **Fig. S1H**). As shown in **Fig. 1D-F**, the contraction area of cardiogenic gastruloids at day 7 can occupy up to 73% of the entire aggregate, but has an average of around 17%. Frequency average in contractile aggregates is at around 0.6Hz and is stable from day 7 to day 10 (see also **Fig. S1F-G**). Finally, tissue deformation per contraction has an average of 7um, but can reach up to 26 um. As expected, aggregates derived from gastruloid protocols displayed no contractility nor deformation. To confirm the contractility area results, we also performed immunohistochemistry staining for the cardiomyocyte marker cardiac Troponin (cTnT). We then measured the area of cTnT+ cells in the aggregates, which, in accordance with the % contracting area, resulted in minimal area for gastruloid protocols at day 7, and an average of 25% for cardiogenic gastruloids (**Fig. S1B-E**). Surprisingly, at day 10, gastruloids also produced a certain amount of cTnT staining (**Fig. S1A**), although this did not result in contraction capacity, possibly indicating that differentiated cells are non-contractile.

As a comparison (although the mechanics of movement are radically different), a *C. elegans* nematode that measures 300-1000μm in length propels itself when muscle contractions elicit a 30-100μm body deformation change; the percentage of the body that is contracting is estimated at 15-20%.

To determine whether cardiogenic gastruloids were motile, we then performed motility assays on the aggregates derived from cardiogenic gastruloid protocols. To do so, we placed 5 to 8 aggregates per experimental set into low-adherence 6-well plates and recorded a timelapse video for 30 minutes using a wide-field microscope at 1X magnification. The samples were placed inside a microscope chamber equipped with an incubator set to 37°C and 5% CO2 for the duration of the experiment. Image tracking software (Imaris) was then used to identify the objects (i.e. the aggregates), and extract their trajectory and total track length over the 30’ period. The total track length is measured as the distance each aggregate traveled from the first image captured at T=0 min to the last image taken 30 minutes later. We first recorded the non-beating aggregates (n=22) generated by the gastruloid protocol after D7 and D10 of differentiation. As expected, the displacement and track length were minimal (under 100um for all aggregates) and there was no global drifting during imaging, demonstrating the stability of the setup. We then measured the track length of cardiogenic gastruloid aggregates: although at day 7 the motility average is non-distinguishable from gastruloids (**Fig. 1H**), a small percentage (2/33) of aggregates already showed a track length above 100um (120um and 378um). On day 10, the difference between gastruloids and cardiogenic gastruloid was more evident (**Fig. 1I**). The average motility for cardiogenic gastruloid protocols was 31um, significantly higher than gastruloids. Moving cardiogenic gastruloids had an average speed ranging from 0.02 to 0.05 µm/s (**Fig. 1J**). Prompted by these results, we also measured shorter-term motility in the 30’’ timelapse movies used for contractility assays (as opposed to the 30’ longer term motility of the motility assay above). For cardiogenic gastruloids the short-term motility is indeed also apparent at day 10, in a range between 0 and 12um, with an average of 3um/min (**Fig. 1G**). For a quantitative comparison, we report in **Fig. S1H** the values for speed of different locomotion types; for example, *C. Elegans*, which has a body plan that is highly optimized for muscle-based motility, moves at 4 orders of magnitude faster at 100–500 μm/s^41,42^.

We then quantitatively assessed which morphology or contractility features of cardiogenic gastruloids were most associated with motility. To do so, we performed mono-variate correlation analysis for the measured contractility and morphology features (see also **Fig. S2**) with track length. As shown in **Fig. 1K**, we observed correlation with track length especially for elongation parameters (eccentricity and aspect ratio), frequency, deformation per contractions, and contracting area percent, whereas having a contraction focus in an eccentric position was found to be negatively correlated with track length. This suggested to us that improving some of these features would potentially improve motility.

Collectively these results show that currently reported cardiogenic gastruloids display contractility features compatible with some minimal motility, at low efficiency. The results also identify some features of the more motile aggregates that are associated with longer track lengths. The next step would be to design developmental trajectories that would lead to aggregates with increased contractility and motility features.

### Novel ‘cardiobot’ protocol generates aggregates with increased cardiomyocytes yields

One challenge with currently available protocols is that they produce contractile aggregates at low efficiency, such that for each differentiation run, only a subset of the aggregates are motile. We hypothesized that deriving protocols with increased efficiency would increase the chance of generating motile aggregates. We designed a small screen where we would modulate the protocol at the mesoderm induction stage, or at the cardiogenic induction phase, and identify the combination of factors that would give rise to the highest % of contracting aggregates at day 7.

For the mesoderm induction phase, cardiogenic gastruloids use Wnt induction with Chiron. We included in the screen induction of mesoderm with Transforming Growth Factor (TGF)-beta family members Activin-A and Bone Morphogenetic Protein 4 (BMP4). In fact, it was previously shown that the right combination of Activin-A and BMP4 promotes the emergence of cardiogenic progenitor cells (CPCs) and ultimately leads to the efficient formation of cardiomyocytes up to 90% ^43^. Activin-A and BMP4 can serve as cardiac induction molecules in other embryoid models, such as cardiac organoids ^44–46^. For the cardiogenic media, applied after mesoderm induction, the cardiogenic gastruloid medium uses N2B27 medium, a mix of neurobasal medium and DMEM/F-12 which is supplemented with N-2, B-27 supplements in addition with soluble factors such as AA, VEGF, FGF2^40^. Another medium that is described in the literature as being more conducive to hematopoiesis and cardiomyocyte development^47^ is StemPro-34 SFM medium. Additionally, we considered addition of FGF10 to the other growth factors as it has been shown to further support cardiac fate development^48^.

Based on these considerations, we devised a screen where mesoderm induction done with one of these variants is followed by cardiogenic induction or basal media, as shown in schematic **Fig. 2A**. We subjected 300 mES cells to these protocols in U-bottom plates, then assessed their contraction efficiency at day 7. As shown in **Fig. 2B**, the combination of Wnt induction and StemPro based cardiomyocyte differentiation medium (CDM) (protocol #2) did not increase the percentage of beating aggregates. On the other hand, the combination of Activin-A/BMP4 (AB) or Activin-A/BMP4/VEGF (ABV) with StemPro-based CDM (protocols #4 and #6) produced contractile aggregates at substantially higher efficiency. We termed protocol #6 as the “cardiobots protocol”.

**Figure 2:**
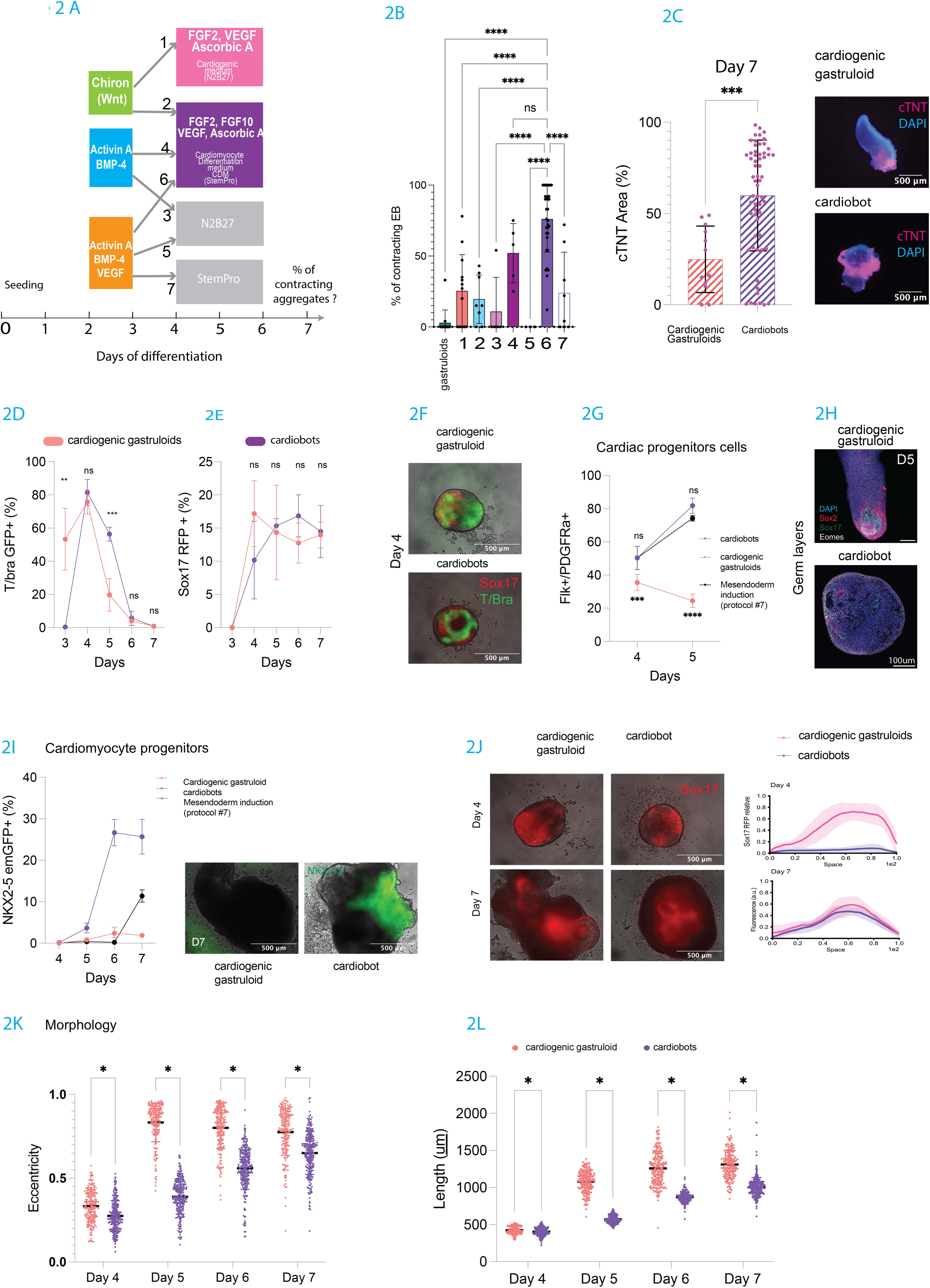
A novel protocol improves differentiation efficiency and enhances cardiomyocyte yield. **A.** Schematic illustration of the screening strategy used to develop various cardiogenic protocols, screening different mesoderm induction and cardiogenic factors. The first induction step (days 2-3 post-aggregation), involved either CHIR99021 (Chiron), a Wnt agonist, or a combination of Activin A and BMP4 with or without VEGF. The second phase (days 4-5) involved the addition of soluble factors such as ascorbic acid, FGF2, and FGF10. Numbers 1-7 indicate the 7 different combinations that were tested (e.g. protocol #1 is obtained using Chiron for mesoderm induction, and AA, FGF, VEGF in N2B27 for cardiogenic induction, see Fig. S2 for list of protocols and names). **B.** Bar graph showing the average percentage of contracting embryoid bodies across different protocols. N = number of independent experiments. Conditions tested: gastruloids (N=13), Cardiogenic gastruloid, protocol #1 (N=13), protocol #2 (N=8), protocol #3 (N=5), protocol #4 (N=5), protocol #5 (N=3), cardiobots, protocol #6 (N=33), protocol #7 (N=8).Statistical analysis was performed using one-way ANOVA. **C.** Left panel: Bar graph showing the percentage of cTnT-positive area on Day 7 for cardiogenic gastruloids (n=12) and cardiobots samples (n=61). Statistical analysis was performed using the t-test. *n* = number of individual aggregates. Right panel: Representative immunofluorescence images of cardiogenic gastruloid and cardiobots samples at Day 7 stained for cardiac troponin T (cTnT, magenta) and DAPI (blue). Scale bar: 500 µm. **D.** Time course of average percentage of cells that express the T-Bra-GFP reporter over the total cells of individual aggregates, cardiogenic gastruloids indicated by red dots, and cardiobots by purple dots. Each dot is an average of N=5 independent experiments, where a minimum n=16 aggregates/per experiment and conditions have been used. Statistic tests have been performed using multiple t-tests. **E.** Time course of average percentage of cells that express the Sox17-mCherry reporter over the total cells of individual aggregates, cardiogenic gastruloids indicated by red dots, and cardiobots by purple dots. Each dot is an average of N=5 independent experiments, where a minimum n=16 aggregates/per experiment and conditions have been used. Statistic tests have been performed using multiple t-tests. **F.** Snapshots of Sox17-mcherry and T/Brachyury-GFP reporters expression in representative individual cardiogenic gastruloid and cardiogenic mesendoderm samples at Day 4. Scale bar: 500 µm. **G.** Time course of average percentage of cells double positive for Flk1 and PDGFRα over the total cells of individual aggregates, cardiogenic gastruloids indicated by red dots, and cardiobots by purple dots. Each dot is an average of N=5 independent experiments, where a minimum n=16 aggregates/per experiment and conditions have been used. Statistic tests have been performed using multiple t-tests. **H.** Immunofluorescence images of representative cardiogenic gastruloid and cardiobot samples at Day 5 stained for markers of germ layers: Sox2 (neuroectoderm, red), Sox17 (endoderm, green), EOMES (pan-mesendoderm, grey), and DAPI (blue). Scale bar: 100 µm.**I.** Time course of average percentage of cells that express the NKX2-5-GFP reporter over the total cells of individual aggregates, cardiogenic gastruloids indicated by red dots, and cardiobots by purple dots. Each dot is an average of N=5 independent experiments, where a minimum n=16 aggregates/per experiment and conditions have been used. Statistic tests have been performed using multiple t-tests.Representative snapshots of cardiobot at Day 7 showing NKX2-5-GFP reporter positivity (green). Scale bar: 1000 µm. **J.** Left panel: Representative images showing Sox17-mCherry reporter (red) in representative cardiogenic gastruloids and cardiobot samples at Day 4 and Day 7. Right panel: Average Sox17-mCherry reporter expression along the length of individual aggregates, red line cardiogenic gastruloids, purple line cardiobots, at day 4 and day 7. **K.** Dot plots showing eccentricity and length measurements for cardiogenic gastruloids (n=195) and cardiogenic mesendoderm samples (n=262) over time. *n* = number of individual aggregates. Averages are indicated with bold lines. Statistical analyses were performed using the t-test and Mann–Whitney test as appropriate. Statistical significance: *p < 0.05; **p < 0.01; ***p < 0.001.

We then focused on this cardiobots protocol and considered whether it would generate aggregates with a higher percentage of cardiomyocytes at day 7 compared to cardiogenic gastruloids, which could mean increased capacity for tissue deformation and motility at later days. To do so, we measured the percentage area of cTnT, and found that the cardiobots protocol generates aggregates with substantially higher percentages of cTnT compared to cardiogenic gastruloids (**Fig. 2C**, cTnT % and **Fig. S2A** cTnT intensity). Thus, it appears that this protocol generates cardiomyocytes more efficiently, both in terms of efficiency per batch, and also in individual aggregates that have larger areas positive for cardiomyocyte markers.

We hypothesized that this increased efficiency of differentiation is due to increased progenitors, and more favored cardiomyocyte differentiation microenvironment. To test these hypotheses, we evaluated the developmental trajectory in the cardiobots protocol and compared it to the cardiogenic gastruloid. We used a dual-reporter mESC line for evaluating in real-time the differentiation trajectories of mesoderm and endoderm lineages (**Fig. 2D-F** and **Fig. S2B** for imaging time course): T-Bra eGFP and Sox17 Strawberry^49^ to monitor the expression of the mesoderm through T-Brachyury, a marker of nascent mesoderm^50^, and Sox17, a marker of definitive endoderm, the main component of the gut tube. We used FACS for measuring the percentage of cells positive for these markers; as shown in **Fig. 2D**, although both protocols show induction of endoderm and mesoderm at day 3-5, they do so with different dynamics. Indeed, T-Bra, is already induced at day 3 in cardiogenic gastruloids, then peaks at day 4, and dips down to around 20% positive cells at day 5. In the cardiobots protocol, T-Bra induction is delayed such that at day 3 it is not yet induced; it still peaks at day 4, but then is significantly sustained (around 60% of cells) at day 5, before subsiding by day 6. By comparing T-Bra in protocol #7, we saw that its expression at day 5 was lower compared to the cardiobots protocol (**Fig. S2C**), suggesting that the CDM induction is influencing sustained T-Bra expression. Early endoderm marker Sox 17 follows a similar trend where the cardiobots protocol induces it at a slightly later time point; but, contrary to T-Bra, it remains sustained throughout the remainder of the differentiation in both protocols (**Fig. 2E**).

We then moved our analysis to cardiomyocyte differentiation. Contractile cardiomyocytes are known to arise from cardiac progenitor cells (CPCs), which give rise to several cell types specific to cardiac development, such as cardiac fibroblasts, endothelial cells, and smooth muscle cells^51^. These CPCs can be detected by the presence of surface markers, namely Flk-1 and PDGFRα^51^. To measure CPC specification, we performed FACS analysis for these markers, and calculated the percentage of precursors at different stages of *in vitro* differentiation at days 4 and 5. The cardiobots protocol generates cardiogenic precursors at a higher percentage compared to cardiogenic gastruloid protocols (**Fig. 2G** Flk-1/PDGFRα double positive: 50.2% vs 35% at day 4, 24.4 vs 81.8% at day 5). Interestingly, it seems that the CDM is again important to maintain elevated CPC in the aggregates: as shown in **Fig. S2D**, protocol #2 for example, where Chiron is followed by CDM, show significantly higher CPCs compared to cardiogenic gastruloids, although still lower than the cardiobot protocols.

The next stage of cardiomyocyte development involves upregulation of NKX2-5, a transcription factor that plays a major role in heart tube formation and directs the differentiation of CPCs into cardiomyocytes^52^. By assessing the expression levels of NKX2-5 during differentiation, we aimed to determine whether the observed differences in cardiomyocyte maturation between cardiogenic gastruloids and the cardiobots protocols could be attributed to variations in NKX2-5 activity. To this end, we used an NKX2-5 emGFP reporter^53^ to monitor the expression of NKX2-5 over time across different protocols. FACS analysis showed that NKX2-5 expression is strongly upregulated in cardiobot aggregates starting from day 6 of differentiation, whereas it was not strongly activated in cardiogenic gastruloids (NKX2-5: 25% vs 2% at day 6). These results were confirmed by time lapse imaging data (**Fig. 2I**, see also **Fig. S2B**,**S2E**).

Of note, ectoderm derivatives are induced in both protocols: by IF at day 5 we observed groups of cells positive for Sox2 in cardiogenic gastruloids, as expected ^38,54^; Sox2 is also induced in the cardiobots protocol, although at lower efficiency (**Fig. 2H** and see **Fig. S2F** for single channels).

We then evaluated the localization and patterning of these nascent germ layers in the aggregates. We confirmed the observation that cardiogenic gastruloids showed asymmetric induction of T-Bra positive cells, marking the initiation of the anterior-posterior axis establishment. In the cardiobots protocol, mesoderm induction with ABV factors results in a more diffuse T/Bra+ cell expression across the aggregate (**Fig. 2F**). Sox17 is largely expressed in cardiogenic gastruloids on the opposite side of where T-Brachyury is expressed from day 4 of differentiation. Later, Sox17 is predominantly expressed in the anterior side of the cardiogenic gastruloid (and all Chiron-based protocols, see **Fig. S2G**) and continues toward the posterior part (where elongation is observed), forming a gut tube-like structure (consistent with what was described previously^33^); in contrast, in the cardiobots protocol, Sox17 expression remained low and started to emerge from day 5 of differentiation onward and was observed in ball-like structures at day 6. Interestingly, at these later days, localization of Sox17 becomes polarized to one area of the aggregate in the cardiobots protocol as well; this, alongside the observation that Nkx2.5 is also patterned asymmetrically, is indicative of a late symmetry breaking event that occurs in the cardiobots protocol at later time points compared to gastruloids protocols (**Fig. 2J** and **S2H**).

Finally, we quantitatively compared simple morphology metrics of the new cardiobots protocol compared to cardiogenic gastruloids (**Fig. 2K-L** and **S2J** for the other protocols). At day 5 the cardiogenic gastruloid aggregates take on the distinctive elongated structure, with high eccentricity (around 0.8); whereas the cardiobots remain more spherical, scoring at around 0.3. By day 7 this difference is much less pronounced (0.5 vs 0.6). Similarly, the area and main length of the aggregates is higher for cardiogenic gastruloids at day 5 and 6, but the cardiobots reduce this difference by day 7. Compared with the other protocols too, it seems indeed that the early elongation morphogenesis is dependent on the mesoderm induction with Wnt stimulation, characteristic of the gastruloids protocols.

Collectively, these results show that the new protocol based on mesoderm induction with ABV factors, and cardiogenesis with CDM, induces mesoderm and endoderm lineages that support enhanced cardiomyocyte differentiation compared to cardiogenic gastruloids, and have a distinctive pattern of morphogenesis.

Our next step is to assess whether we can control the location of the cardiomyocytes as this contraction is essential for generating the force needed to induce movement within the entire aggregate.

### Synthetic organizers can be used to control localization of cardiogenic areas in cardiobot protocols

Use of synthetic organizers have been described to control localization of differentiation in stem-cell derived organoids^28^ [add kidney ref] and in particular for controlling cardiomyocyte localization in embryoid bodies with Wnt induction^27^. We sought to test the hypothesis that emergence of localized signaling hubs could give rise to contractile, motile biobots with defined morphologies and contractile areas. To test this hypothesis, we mimicked localized hubs through synthetic organizers. We genetically engineered HEK cells with doxycycline-inducible cassettes for the expression of mesoderm inducers Wnt3a or BMP4 coupled to BFP (induction reporter), alongside constitutive Cdh3 and mCherry expression for enhanced cohesion and tracking. We also generated HEK cells with dox-inducible cassettes for secretion of cardiogenic factors FGF2, FGF10 and VEGF, alongside a GFP induction reporter and Cdh3 for cohesion. We characterized the expression of inducible cassettes to increasing dox concentrations via flow cytometry and dose-response curves (EC50=2.61nM +/−0.863nM across cell lines) (**Fig. S3A-C**).

We then set out to investigate the developmental consequences of conjugation of spheroids of HEK cells with spheroids of mES cells. To do so we pre-aggregated 500 engineered HEK cells into Spheroids using U-bottom plates for 2 days prior to the co-culture with or without Dox. Individual HEK spheroids were then moved into single wells where embryoid bodies are grown at different stages of the differentiation trajectory. We term this association a hybrid assembloid.

We first assessed mesoderm induction in the first phases of differentiation by means of Wnt3a and Bmp4 organizers. As shown in **Fig. 3A-C**, embryoid bodies co-cultured with CDH3 spheroids, lacking a drug-inducible morphogen cassette, showed no induction of nascent mesoderm/primitive streak reporter T-Bra. In contrast, Wnt3a and BMP4 organizers induced T-Bra fluorescence at day 4, to levels similar to cardiobots and gastruloids; BMP4 organizers induced the highest intensity of T-Bra fluorescence (**Fig.3B**). Moreover, T-Bra+ cells tended to localize towards the organizer especially for BMP4 organizers (**Fig.3C**). Of note, Wnt3a organizer-conjugated embryoid bodies elongated on day 5 of culture, similar to the CHIR condition (gastruloids/cardiogenic gastruloids), without provision of additional factors in the basal media (N2B27) (**Fig. S3D-F**). Altogether, these results suggest successful mesoderm induction triggered by Wnt3a and BMP4 organizers in the embryoid bodies.

**Figure 3:**
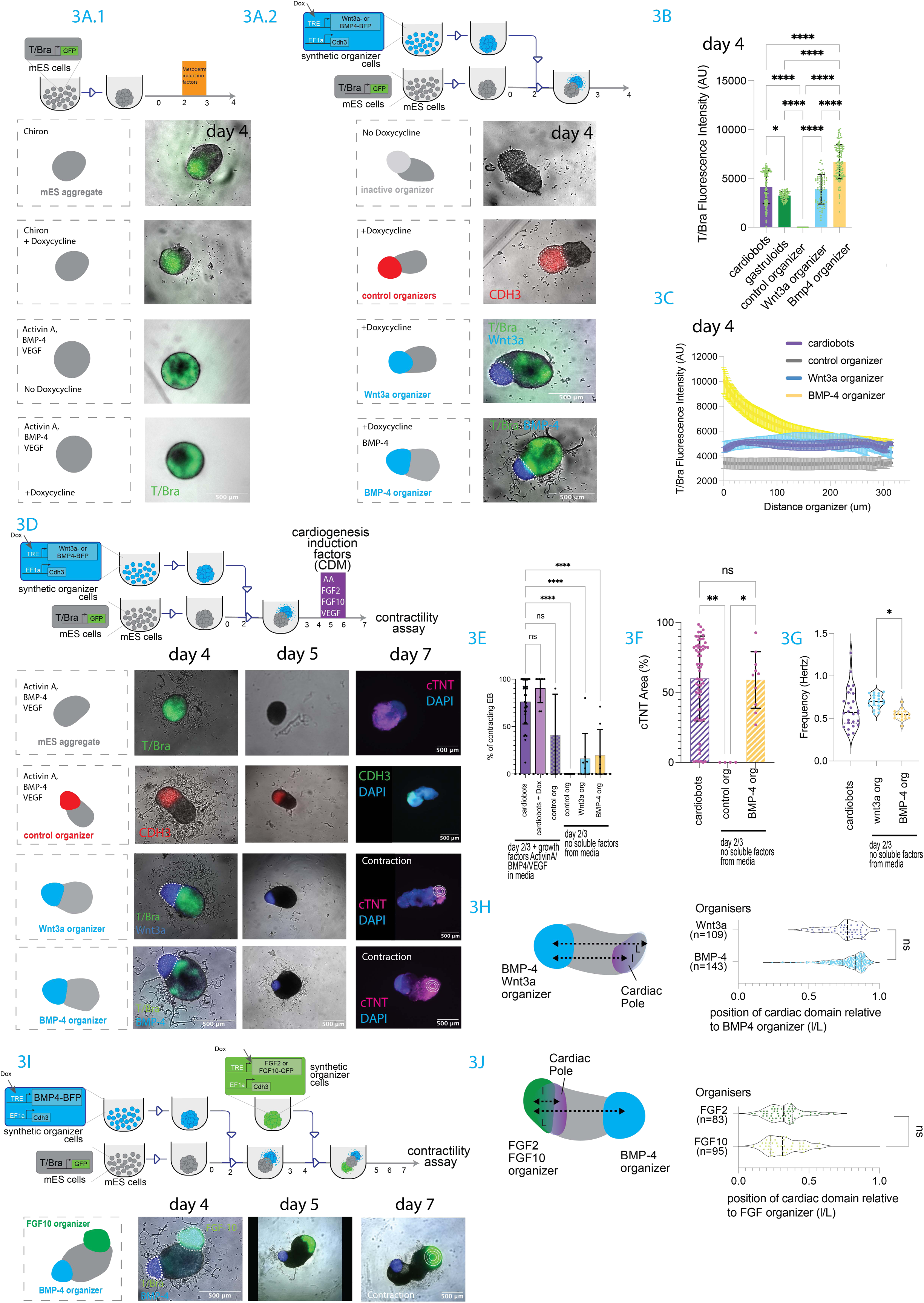
Synthetic organizers can induce cardiogenic fate in embryoid bodies. **A.1.** Top, schematics of experiment, with mES T-Bra-GFP reporter cell aggregation, and timeline of factors stimulation on the right. Bottom: Representative overlay of microscopy picture with bright field and green fluorescent channel of assembloids at Day 4. Scale bar 500um. **A.2** Top, schematics of experiment, with mES T-Bra-GFP reporter cell aggregation, in parallel to synthetic organizer cell aggregation; conjugation of pre-aggregated spheroids at day 2, monitoring till day 4, no soluble factors addition. Bottom, representative overlay of microscopy picture with bright field and green fluorescent channel of assembloids at Day 4. cell bodies are shown in grey, and BMP-4 and Wnt3a organizers are shown in blue (BFP reporter), control organizer is shown in red (mCherry reporter). The nascent mesoderm marker T/Brachyury (T-Bra) is shown in green. **B.** Bar graph showing T--GFP fluorescence intensity (arbitrary units based on image analysis of fluorescent microscope images, see methods) across different conditions. *n* = number of individual aggregates: cardiobots (n=138), gastruloids (n=79), assembloid with control organizer (n=77), Wnt3a organizer (n=66), BMP4 organizer (n=134). Statistical analysis was performed using the Kruskal–Wallis test. **C.** Line plots showing T-Bra-GFP intensity as a function of distance from the organizer (in µm) for each condition. Purple line: cardiobots (positive control), grey line: control organizer (negative control), blue line: Wnt3a organizer, yellow line: BMP4 organizer. **D.** Top, schematics of experiment, with mES T-Bra-GFP reporter cell aggregation, in parallel to synthetic organizer cell aggregation; conjugation of pre-aggregated spheroids at day 2, addition of cardiogenetic factor in the media between days 4-6. Bottom, time-course snapshots of assembloids combined with the indicated organizers (or no organizer). Concentric white circles indicate contraction zones. Organizers (Wnt3a and BMP4) are shown in blue. The control organizer in the second row is shown in red. T/Brachyury-GFP expression is in green. Scale bar: 500 µm. **E.** Bar graph showing the average percentage of contracting assembloids for each condition at Day 7. Conditions: mesoderm induction done with factors: cardiobots (N=33), cardiobots + Doxycycline (N=9), ABV + control organizer (N=3); mesoderm induction done with synthetic organizers: control organizer (N=8), Wnt3a organizer (N=8), and BMP4 organizer (N=8). Statistical analysis was performed using ANOVA. **F.** Bar graph showing the percentage of cTnT-positive area per assembloid on Day 7. Conditions: cardiobots (N=39), control organizer (N=4), BMP4 organizer (N=10). Kruskal–Wallis test was used for statistical analysis. **G.** Violin plots showing contraction frequency (in Hz) in assembloids. Black dotted line represents the median. *n* = number of individual aggregates: cardiobots (purple, n=26), Wnt3a organizer (blue, n=20), BMP4 organizer (yellow, n=15). **H.** Violin plots showing the relative distance between organizers (BMP4, n=143 or Wnt3a, n=109) and contraction zones. *l*: distance from the organizer center to the center of the contraction zone *L*: distance from the organizer center to the edge of the entire aggregate. Statistical analysis was performed using the Mann–Whitney test. **I.** Top, schematics of experiment, with mES T-Bra-GFP reporter cell aggregation, in parallel to mesoderm induction synthetic organizer cell aggregation; conjugation of pre-aggregated spheroids at day 2, addition of cardiogenetic synthetic organizers at day 4. Bottom, time-course snapshots of assembloids combined with the Bmp4 mesodermal induction organizer, and FGF10 cardiogenic organizer. Concentric white circles indicate contraction zones. BMP4 organizer is shown in blue. The FGF10 organizer is shown in green. Scale bar: 500 µm. **J.** Violin plots showing the relative distance between cardiogenic organizers FGF2 (n=83), FGF10 (n=95) and contraction zones. *l*: distance from the organizer center to the center of the contraction zone. *L*: distance from the organizer center to the edge of the other side of the aggregate.Statistical analysis was performed using the Mann–Whitney test. Statistical significance: *p < 0.05; **p < 0.01; ***p < 0.001.

We then assessed if mesoderm cells generated via synthetic organizers could be driven to cardiac fates. To do so, we continued assembloid culture in presence of soluble cardiogenic factors in culture media. Remarkably, when assembloids containing BMP4 or Wnt3a organizers were treated with cardiogenic soluble factors at day 5, they gave rise to contractile embryoid bodies at Day 7 (**Fig. 3D-G**). Wnt3a and BMP4 organizers supported generation of contractile aggregates with efficiency around around 15-20% on average (**Fig. 3D-E**). The cardiogenic areas of the BMP4 organizer’s aggregates were comparable to the soluble-factors protocol, at an average of 58%, indicating a robust induction of large areas of contractile cells (**Fig. 3D, F**).

Contraction frequency was slightly higher for Wnt3a organizers compared to BMP4 organizers (**Fig. 3G**); BMP4 organizers had an average contraction frequency that most closely mimicked the cardiobots protocols with soluble factors. Of note, organizers that are engineered to secrete other cardiobots mesoderm induction factors such as VEGF and Activin-A are unable on themselves to support generation of contractile aggregates (**Fig. S4B**).

As a control, we run cardiobots protocols with control organizers: here the % of contractile cardiobots reduced to an average of 40 % by day 7 compared to cardiobots protocol without organizers (above 75%), indicating reduced efficiency of cardiogenic differentiation in the presence of a control organizer (**Fig. 3E** and also **Fig. S4A**). This made the results of cardiogenic induction with BMP4 and Wnt3a organizer even more remarkable.

We then asked whether localization of the contractile area was influenced by the organizers: as shown in **Fig. 3H**, the localization of the contractile domains is to the opposite side of the organizer. The distance of contractile domains from the organizers is on average 77% of the entire length of the aggregate for Wnt3a organizers and 78% for BMP4 organizers. This indicates that the presence of organizers skew cardiogenic fates to appear away from the organizer themselves.

This shows that mesoderm induction with synthetic organisers secreting Wnt3a or BMP4 can support cardiogenesis with CDM soluble factors, and that the localization of the contractility is always on the opposite side of the organizers.

We finally asked if the cardiogenesis induction phase could also be replaced by organizers. We reasoned that this would give rise to a construct where the contractions were proximal to the CDM organizers, and that this would be generated without addition of soluble factors. To do so, we generated organizers that produce CDM factors FGF2 and FGF10, then we conjugated first BMP4 organizers on day 2 and then FGF organizers (either FGF2 or FGF10) on day 4 to embryoid bodies in basal medium (StemPro). We then screened the aggregates where the 2 organizers took diametrically opposing attachment sites (approximately 50% of the total), and calculated the position of the contractile areas relative to the FGF organizers. As shown in **Fig. 3I-J**, the localization of contractile areas was proximal to the FGF organizers (around 30% the length of the entire aggregate) (see also **Fig. S4C**).

These results show that for cardiobots, emergence of a localized mesoderm-inducing signaling hub is sufficient to position the emergence/location of a contractile domain precursor. Maturation of the cardiac domain precursor into a functional beating population requires input in the FGF pathway. This suggests that self-organization of a contractile apparatus would need coordination with at least two signaling hubs (or rewiring of the precursor domain motility, so that it displaces itself from the mesoderm hub towards an FGF hub).

Altogether, **Fig. 2** and **Fig. 3** showed that the novel cardiobots protocol, based on ABV mesoderm induction, and CDM cardiogenesis, is modulable, and can give rise to aggregates with high percentage of cardiomyocytes across 7 days of differentiation.

### Cardiobots develop contractility and motility features

To assess motility capabilities of the cardiobots protocols and compare them to cardiogenic gastruloids (**Fig. 4A** for compared protocols), we proceeded with motility assays on the cardiobots. As shown in **Fig. 4B**, at day 7, when the earliest cardiomyocyte contractions are typically observed, motility is low across all the protocols, with averages below 100 um. We did notice that cardiobots show a larger group of aggregates with high motility. We then compared motility at day 10 of differentiation (**Fig. 4C** and S5L for more examples), when the cardiogenic gastruloid showed increased motility (See **Fig. 1I**). For cardiobots, we compared two conditions: one where aggregates are exposed to cardiogenic media CDM only between day 4 and 6 (we call this the cardiobots protocol); and another where cardiogenic media CDM exposure is continued throughout day 10 (the extended cardiobots protocol, **Fig. 4A**). As shown in Fig. 4D, both cardiobots and extended cardiobots displayed significantly increased motility compared to cardiogenic gastruloids; as shown in Fig. 1, cardiogenic gastruloids have average track length in the same assay of 31um, and none of them have motility higher than 100um. In contrast, the average motility for cardiobots was 367 μm (with a maximum of 2733 μm), and even more striking it was 1060 μm for extended cardiobots (maximum of 7283μm). This results in values of speed that are around 10x faster than cardiogenic gastruloids: Moving cardiogenic gastruloids had an average speed ranging from 0.002 to 0.05 µm/s, whereas cardiobots have speed of 0.2 μm/sec, and extended cardiobots of 0.8 μm/sec. Moreover, 40% of the cardiobots (34 out of 83) and 80% of the extended cardiobots (66 out of 81) exhibited track lengths greater than 100 μm. Similar trends were obtained for measures of micromotility (**Fig. S5K**) and displacement (i.e. distance calculated from initial position to final position after 30’ timelapse), indicating that circular rotation in place is a minority of the trajectories (**Fig. S5A,B**). These results indicate that motile cardiobots can be generated with Activin/BMP4/VEGF mesoderm induction, followed by cardiogenic media. These cardiobots are consistently more motile compared to cardiogenic gastruloids.

**Figure 4:**
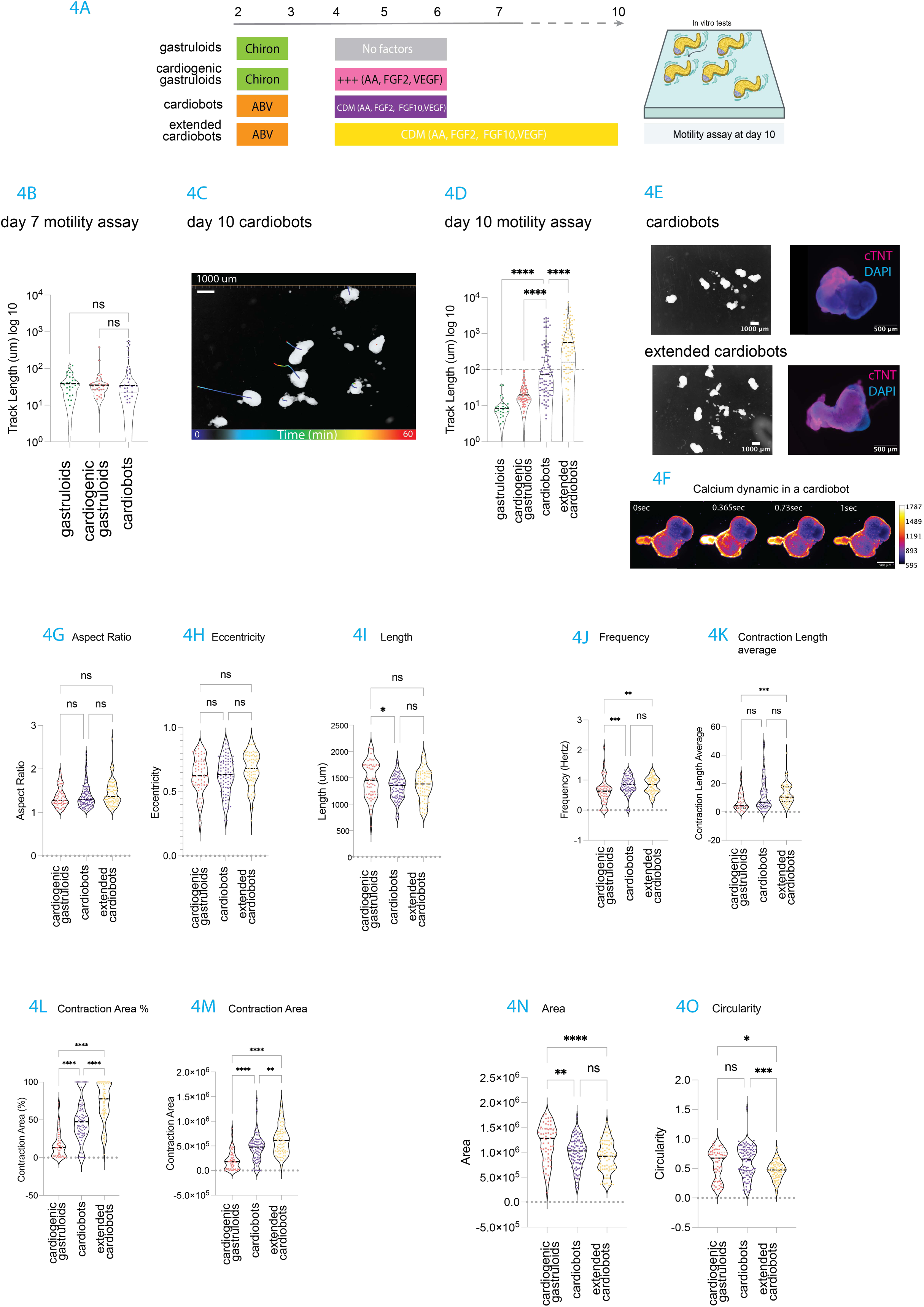
Novel cardiogenic embryoid bodies display enhanced motility. **A.** Schematic illustration summarizing the protocols used to differentiate stem cells into various cardiogenic embryoid bodies (EBs) up to Day 10 of culture. Subsequently, all EBs were assessed using motility assays and feature extraction. **B.** Violin plots showing track length at Day 7 for gastruloids (green, *n* = 27), cardiogenic gastruloids (red, *n* = 29), and cardiobots (purple, *n* = 33). Black dotted line indicates the median. *n* refers to the number of individual aggregates. Track length was recorded over 30 minutes. Statistical analysis was performed using the Kruskal–Wallis test. **C.** Representative snapshots from motility assays at Day 10. Track paths are color-coded using a rainbow scale. Time captions indicate a 60-minute recording. Scale bar: 1000 µm. See also Fig. S5L for other examples. **D.** Violin plots showing track length at Day 10 for gastruloids (green, *n* = 22), cardiogenic gastruloids (red, *n* = 77), cardiobots (purple, *n* = 83), and extended cardiobots (yellow, *n* = 81). Track length was recorded over 30 minutes. Black dotted lines indicate the median. Gray dotted line at 100um indicates threshold for motility. Statistical analysis was performed using the Kruskal–Wallis test. **E.** Left: Snapshots from motility assays at Day 10 for cardiobots (short and extended protocols). Scale bars: 1000 µm. Right: Fluorescent microscope image of individual aggregates stained with cTnT (purple) and DAPI (blue) at Day 10; scale bar 500 µm. **F.** Time-course snapshots showing calcium dynamics in a representative cardiobot. Scale bar: 500 µm. Intensity of calcium signal is coded with color intensity as represented in the legend on the right. **G-O**. Violin plots for the indicated features for aggregates at day 10; black dotted line indicates the median. Statistical analysis was performed using the Kruskal–Wallis test. For description of how the features are calculated, see Methods. **G.** Morphological aspect ratio of aggregates at Day 10 for cardiogenic gastruloids (red, *n* = 53), cardiobots (purple, *n* = 88), and extended cardiobots (yellow, *n* = 73). **H.** Morphological eccentricity of aggregates at Day 10 for cardiogenic gastruloids (red, *n* = 53), cardiobots (purple, *n* = 88), and extended cardiobots (yellow, *n* = 73). **I.** Aggregate length (µm) at Day 10 for cardiogenic gastruloids (red, *n* = 49), cardiobots (purple, *n* = 87), and extended cardiobots (yellow, *n* = 73). **J.** Contraction frequency (Hz) at Day 10 for cardiogenic gastruloids (red, *n* = 56), cardiobots (purple, *n* = 82), and extended cardiobots (yellow, *n* = 74). **K.** Average contraction length (µm) at Day 10 for cardiogenic gastruloids (red, *n* = 35), cardiobots (purple, *n* = 73), and extended cardiobots (yellow, *n* = 62). **L.** Contraction area as a percentage of total aggregate area at Day 10 for cardiogenic gastruloids (red, *n* = 56), cardiobots (purple, *n* = 82), and extended cardiobots (yellow, *n* = 74). **M.** Contraction area (µm²) at Day 10 for cardiogenic gastruloids (red, *n* = 52), cardiobots (purple, *n* = 86), and extended cardiobots (yellow, *n* = 70). **N.** Aggregates area (µm²) at Day 10 for cardiogenic gastruloids (red, *n* = 53), cardiobots (purple, *n* = 88), and extended cardiobots (yellow, *n* = 73). **O.** Aggregates circularity at Day 10 for cardiogenic gastruloids (red, *n* = 53), cardiobots (purple, *n* = 88), and extended cardiobots (yellow, *n* = 73). Statistical significance: *p < 0.05; **p < 0.01; ***p < 0.001.

What are the contractility and morphology features of these cardiobots? We first performed immunostaining for cardiomyocytes in cardiobots, and saw that cTnT+ cells continue to be present at day 10 (compare with results at day 7 in **Fig. 2C**), and seem to be more widespread in extended cardiobots (**Fig. 4E**, see also **Fig. S5C-F** for quantifications). We also performed calcium assays (**Fig. 4F**) as described above, to capture frequency and other contractility features. We looked at the contractile and morphological features, starting from the ones that were correlated with motility in cardiogenic gastruloids (see **Fig.1K**). Surprisingly, for many of the highly correlated features like aspect ratio, eccentricity and length (**Fig. 4G-I**), we measured limited to no difference between cardiogenic gastruloids and cardiobots. Small but statistically significant differences could be instead observed for other positively correlating features such as frequency and contraction length average (**Fig. 4J,K**). Frequency for example, was higher in the cardiobots protocols compared to cardiogenic gastruloids (0.64 vs 0.83 Hz respectively, see also **Fig. S5G-J**). Of the positively correlating features however, it was contraction area and contraction area percent that showed the most pronounced increase in cardiobots protocols: contraction area percent for example is at an average of 16.94% for cardiogenic gastruloids, but increases to 49.19% for cardiobots, and to 72% for extended cardiobots (**Fig. 4L,M**). Of the negatively correlated features, circularity and area both show a decrease in the cardiobots protocols (**Fig. 4N,O**). These results show that motile cardiobots have distinctive morphological and contractile properties compared to less motile counterparts at day 10 of differentiation. Moreover, they suggest that the cardiobots have specific features that distinguish them from cardiogenic gastruloids, and prompt the question of which of these features are supporting their increased motility.

### Cardiobots motility depends on a complex relationship between shape, contraction area and frequency

Finally, to identify patterns/parameters implicated in cardiobot motility, we performed mono and multi-variate correlative analysis of cardiobots features with their motility. As for most of Fig. 4, we focused on the cardiobots, extended cardiobots, and cardiogenic gastruloid protocols. We first calculated a correlation matrix for the measured features (**Fig. S6A**) with each other, considering all the aggregates generated from all the 3 protocols together. Although there are areas of high cross-correlation, most of the features are sufficiently independent from each other, with the exception of AR and eccentricity, and area and length, which are natively highly correlated features. We focused on the correlation of morphological and contractility features with the motility measure (track length) (**Fig. 5A**). We noticed that all features’ correlative values were below 0.4, and that contractility features (purple dot) are more highly correlated with motility, with the exception of contraction eccentricity which was anti-correlated. The morphological features were instead either non-correlating, or weakly anti-correlating with motility. This gave a ranked idea of what were the features of the cardiobots that, individually, correlated with the total track length the most. This result is in line with the average observations of **Fig. 4G-O** that protocols that generate more highly motile aggregates (cardiobots and extended cardiobots) have higher contracting area percent and contracting area. Given the variability of features, we also run the monovariate correlation analysis with track length including a feature that indicates the experiment id: although for cardiobots protocol, experiment ID does not correlate with track length, it does so in extended cardiobots at 0.19 correlation value (**Fig. S6C**).

**Figure 5.**
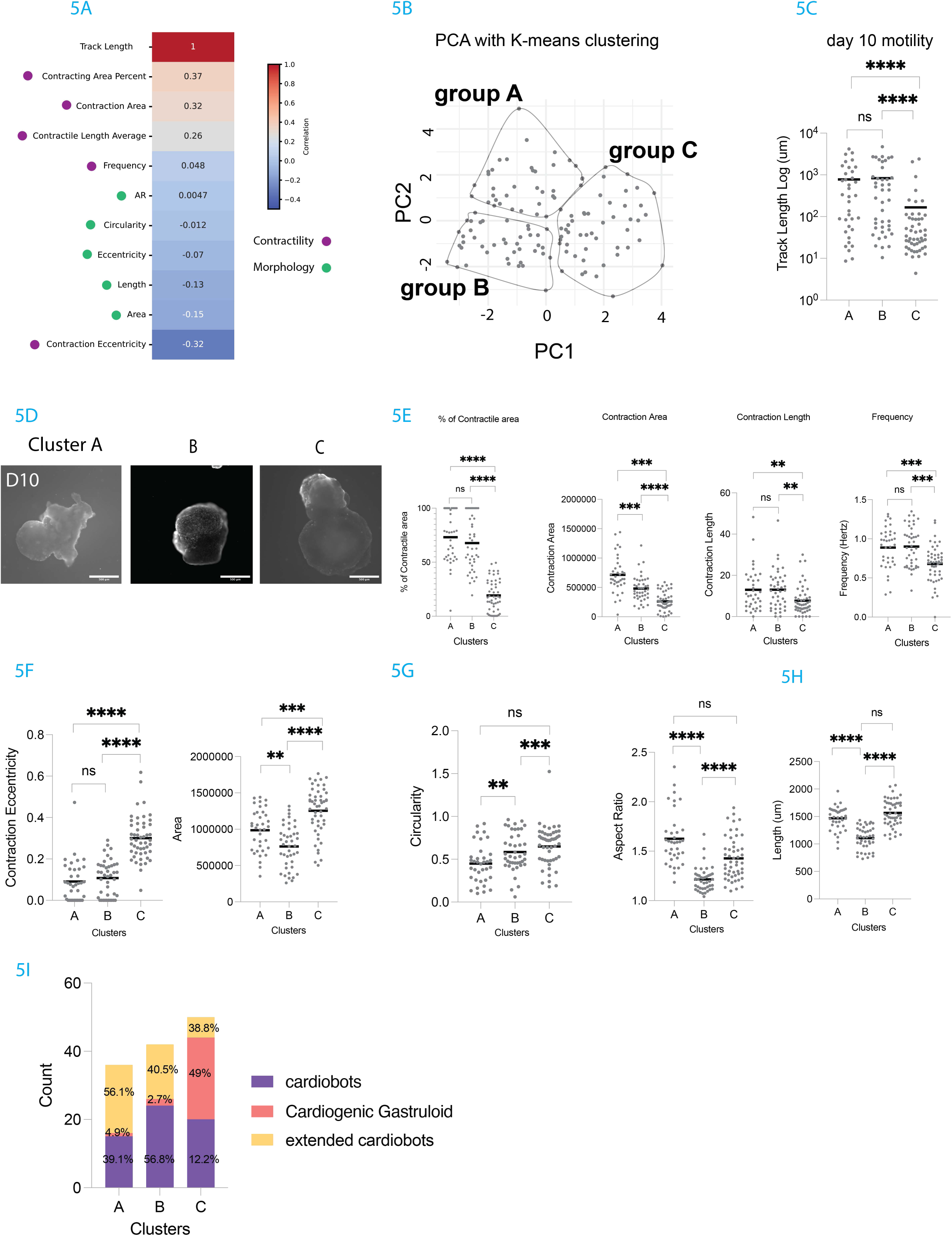
: Classification of motile aggregate. **A.** Correlation matrix for cardiobots (n=59), extended cardiobots (n= 42)and cardiogenic gastruloids (*n* = 3), illustrating the relationships between track length and the indicated morphological (green dot) and contractile (purple dot) features. *n* = number of individual aggregates. The strength of correlation is indicated by the numbers, and by the shade of color, as shown in the correlation legend on the right. **B.** Principal Component Analysis (PCA) plot with K-means clustering performed on all embryoid bodies (*n* = 128), revealing classification into three clusters: Cluster A (*n* = 36), Cluster B (*n* = 42), and Cluster C (*n* = 50). See also Table 1 and 2 for PCA weights, and methods for details. **C.** Dot plot showing track length (μm, log₁₀ scale) for each cluster. Individual aggregates are shown, with cluster averages indicated by bold horizontal lines. Statistical analysis was performed using the Kruskal–Wallis test. **D.** Representative microscope picture of embryoid bodies from each identified cluster (A, B, and C). Images illustrate morphological differences corresponding to clustering analysis. Scale bar: 500 μm. **E.** Dot plot graphs for % contractile area, contraction area (μm²), contraction length (μm) and frequency (Hz), as indicated. Dots are individual aggregates, with cluster averages indicated by bold lines. Statistical analysis was performed using the Kruskal–Wallis test. **F.** Dot plot graphs of contraction eccentricity and area of aggregates (μm²). Dots are individual aggregates, with cluster averages indicated by bold lines. Statistical analysis was performed using the Kruskal–Wallis test. **G.** Dot plot graphs of circularity and aspect ratio. Dots are individual aggregates, with cluster averages indicated by bold lines. Statistical analysis was performed using the Kruskal–Wallis test. **H.** Dot plot graphs of length of aggregates. Dots are individual aggregates, with cluster averages indicated by bold lines. Statistical analysis was performed using the Kruskal–Wallis test. **I.** Stacked bar graph showing the contribution of the aggregates generated with different protocols to the 3 identified clusters. Statistical significance: *p < 0.05; **p < 0.01; ***p < 0.001.

**Table 1:** Principal components weights for the corresponding features shown on the first column. For PC1 and PC2, green highlights positively correlated features above 0.35, and red negatively correlated features under −0.25.

|  | PC1 | PC2 | PC3 | PC4 | PC5 | PC6 | PC7 | PC8 | PC9 | PC10 |
| --- | --- | --- | --- | --- | --- | --- | --- | --- | --- | --- |
| Area | 0.3914317 | 0.11640344 | 0.49114093 | 0.239974451 | -0.04958149 | -0.10521419 | 0.2625598 | 0.387153274 | -0.438702411 | -0.33107525 |
| AR | 0.1690906 | 0.55025832 | -0.30788937 | -0.128381057 | 0.18210393 | 0.11447169 | 0.06011339 | -0.515129307 | -0.466413724 | -0.15673984 |
| Circularity | 0.1333033 | -0.18335859 | 0.42754621 | -0.255860596 | 0.66524495 | 0.48933314 | -0.04085784 | -0.091466289 | 0.083357731 | 0.0369827 |
| Length | 0.3979296 | 0.32470237 | 0.298928 | 0.20497741 | -0.16077488 | -0.09725338 | 0.07404405 | -0.394921898 | 0.63857947 | -0.01761813 |
| Eccentricity | 0.1906437 | 0.54285003 | -0.27708266 | -0.17758354 | 0.16776583 | 0.13040542 | -0.06509516 | 0.644409754 | 0.264509431 | 0.161371 |
| Frequency | -0.2294947 | 0.08005228 | -0.08628775 | 0.522066328 | 0.66067237 | -0.4643629 | 0.06299834 | 0.003001691 | 0.065894272 | 0.02200522 |
| Contraction_Area | -0.2988765 | 0.36662076 | 0.4350642 | 0.213159119 | -0.13990633 | 0.10198958 | -0.42720267 | -0.030138791 | -0.246788023 | 0.51696607 |
| Contracting_Area_Percent | -0.4854991 | 0.23496605 | 0.13390448 | -0.002088963 | -0.03331058 | 0.15937776 | -0.28133788 | 0.059608185 | 0.18601143 | -0.73981858 |
| Contraction_Eccentricity | 0.4610219 | -0.19231375 | -0.11092811 | 0.025009772 | 0.07469453 | -0.23291483 | -0.80620967 | -0.001287868 | -0.073701305 | -0.14935718 |
| Contractile_Length_Average | -0.1244236 | 0.13005804 | 0.29680687 | -0.684320361 | 0.07130512 | -0.63579095 | 0.02152949 | -0.026242503 | -0.004735441 | 0.02694139 |

Do particular combinations of variables are co-correlating with increased motility in individual cardiobots? To get insight to this question, we performed principal component analysis (PCA) on the contractility and morphology features alone, without providing the motility values, for n=129 of total aggregates. Although the data does not strongly cluster immediately, as shown in **Fig. 5B**, k-means clustering (elbow method) identified 3 groups. We asked if these clusters, identified based on contractility and morphology measures, display differences in total track length. We plotted total track length in each cluster: as shown in **Fig. 5C**, total track length for clusters A and B is significantly higher than that for cluster C. **Fig. 5D** shows representative aggregates from the 3 clusters. Similar trends are observed for the displacement length (**Fig. S6B**). The multivariate analysis allows us to identify principal components that distinguish clusters A and B: they are characterized by low values of principal component 1 (PC1). As can be seen for the weights of the principal components decomposition, (**Table 1**), PC1 weighs negatively area and contraction eccentricity, and weighs positively contraction area and contraction area percent. Particularly motile aggregates in clusters A and B are characterized by low area, low length, low contraction eccentricity; and high contraction area and contraction area percent. These are combined features that characterize more motile clusters A and B compared to cluster C.

What distinguishes clusters A and B is instead principal component 2 (PC2). PC2 is especially weighing morphology features related to elongation (aspect ratio and eccentricity for example, see Table 2), such that aggregates with higher values of PC2 indicate more elongated aggregates. Cluster B is lower on PC2, hence representing a cluster of aggregates that are more spherical; whereas cluster A have more elongated aggregates. Since both these classes have similar motility features, it is possible that the elongation features are not particularly relevant for motility; another alternative explanation would be that there are 2 distinct modalities of motility that have similar motility output, but with different mechanical implementation, one for elongated structures, and one for more spherical ones.

**Table 2:** Cumulative proportion of explanation of variance for the indicated principal components.

| Component | Standard Deviation | Proportion of Variance | Cumulative Proportion |
| --- | --- | --- | --- |
| PC1 | 1.8181 | 0.3305 | 0.3305 |
| PC2 | 1.4952 | 0.2236 | 0.5541 |
| PC3 | 1.1958 | 0.143 | 0.6971 |
| PC4 | 1.0272 | 0.1055 | 0.8026 |
| PC5 | 0.90725 | 0.08231 | 0.88489 |
| PC6 | 0.842 | 0.0709 | 0.9558 |
| PC7 | 0.48334 | 0.02336 | 0.97915 |
| PC8 | 0.29368 | 0.00862 | 0.98778 |
| PC9 | 0.25287 | 0.00639 | 0.99417 |
| PC10 | 0.24143 | 0.00583 | 1 |

Overall, this initial analysis showed that morphological and contractility features can generate, on average, distinguishing metrics for cardiobots motility. Consequently, when we plotted the percentage contraction area, contraction area, contractile length average per cluster and frequency, the readings for cluster A and B were significantly higher compared to the ones in cluster C (**Fig. 5E**); on the other hand, contraction eccentricity and area were significantly higher in cluster C compared to cluster A or B (**Fig. 5F**). Moreover, morphology metrics for circularity are higher in cluster B compared to cluster A (**Fig. 5G**), and overall B cluster aggregates are smaller than A cluster aggregates (**Fig. 5H**).

Do the different protocols preferentially produce aggregates that belong to a specific cluster? To answer that question we plotted the counts of aggregates coming from the specific protocol in the 3 clusters: as shown in Fig. 5I, both cardiobots protocols equally generate aggregates that belong to the clusters A and B, whereas cardiogenic gastruloids predominantly generate aggregates that are classified in cluster C. This shows that protocols can skew morphological and contractility properties that are linked to motility.

All in all, this analysis showed that cardiobots with high contractility area, able to generate elevated tissue displacement per contraction, which have non-eccentric contractions and have lower area, are more likely to be motile, and are generated at high efficiency with the new cardiobots protocols.

## Discussion

Controlling developmental trajectories of stem cells in vitro is a frontier of two main efforts: organoids and soft bio-robots. Both of these efforts are discovering to what extent stem cells, with their endogenous genomes, can be influenced in their capacity of self-organization towards user-desired outcomes. Although organoids are showing that stem cells can be driven to generate tissue constructs reminiscent of their natural counterparts, the derivation of self-constructing biobots unconstrained by the guide of in vivo organs, has shown remarkable plasticity of the macro-scale rules of self-assembly ^55,56^. Self-motile, fully-organic biobots have been demonstrated with frog cells and with human cells ^30,32^. The biobots generated with self-organization protocols in these studies were motile thanks to cilia movement. Several studies in the soft robotic field showed that motile biobots can be generated using contractile cells (like skeletal or cardiac muscles) when combined with a supportive scaffold ^9,57,58^. But so far none of the muscle-propelled biobots have been generated via a developmental trajectory. We show here that mouse embryonic stem cells can be driven to follow a continuous developmental trajectory that leads to the formation of aggregates with high contractility that support contraction-propelled motility, which we call cardiobots.

We showed that currently available cardiogenic protocols^40^ and modifications^27^, being geared towards recapitulation of mouse gastrulation and embryogenesis, generate limited contractile tissue. We find here that, although in some isolated cases these can result in motility capacity acquisition at later time points, most of the aggregates generated with these protocols don’t develop strong enough contractile forces to move. This makes sense as, in modern embryos, cardiac contractions support blood flow and not organism movement. In order to generate contractile aggregates we had to increase the contractile areas. To do so, we deconstructed current cardiogenic organoid protocols in two steps: mesoderm induction, and cardiogenic induction, and screened factors used for each of these stages in isolation or in combination. The most optimal protocol we identified involves mesoderm induction with Activin/BMP4/VEGF, and cardiogenesis with FGF2/10/VEGF. Although in gastruloids Activin is usually associated with induction of endoderm, and BMP4 with induction of mesoderm^33^, in 2D differentiation protocols to generate cardiomyocytes from mES cells, BMP4 and Activin have been used in combination to achieve high efficiency of contractile cells^47^. In our cardiobots protocol, Nkx2.5 expression is present in up to 30% of cells by day 7 and, by day 10, aggregates can contain contraction areas estimated at 100% coverage of the cardiobots. These results show that the developmental potential of mES cells is plastic to expansion or contraction of germ layer populations based on cocktails of factors, possibly laying the foundation for screenings based on desired functional outcomes, not necessarily the endogenous ones. There will likely be constraints on how much freedom of operation there is in terms of body plans with only soluble factors. Other strategies, such as synthetic genetic circuits, or synthetic organizers, may be needed to increase control to completely user-defined structures.

Although optimized protocols generate cardiobots with high contraction capacity, they do not provide capacity to control localization of the contractile area in the ‘body plan’ arrangement. Recently, synthetic organizers have been proposed and used to control localization of morphogenetic fields in stem cell derived structures (39706189). We showed here that the cardiobots protocols can be recapitulated, at least in part, with factors provided by synthetic organizers, i.e. engineered self-adherent cell clusters that secrete BMP4 and FGF factors. In these cases, the localization of the contractile areas can be controlled in relationship with the localization of the organizer clusters: on the opposite pole of BMP4 organizers, or in the vicinity of FGF-organizers. It is possible that, by extending the synthetic organizer technology to higher refinement of spatial control, more complex body plans can be achieved. Although synthetic organizers allowed controlled localization of contractile areas, cardiobots generated with synthetic organizers were not able to generate enough contraction to support motility. One challenge with this synthetic organizer approach was that, although designed to be temporally controlled with small molecule induction, the ‘switch off’ phase proved problematic. Combined with the fact that organizers themselves may contribute unwanted additional load, one solution would be a kill switch that may enable transient niche signals, like in the growth-factor-based protocol, and to remove the additional load.

We showed that our aggregates with the cardiobots and even more so with the extended cardiobots protocol have contractions that lead to motility. Motility is dependent on the contractions, as non-contracting aggregates are never motile. We suspect that this kind of motility is reminiscent of burrowing or slithering^59^. As a numerical ballpark comparison (although the mechanics of movement may be radically different), in the nematode *C. Elegans* the amplitude of body deformation required to generate thrust is around 30–100 μm for animals 300–1000 μm long^60,61^; in cardiobots, deformation due to contractions can reach a maximum of 40um, compatible with the highly evolved body plan of C.Elegans. Unlike the C.Elegans example, where the % of its volume that is contracting can be estimated to 15-20% ^62–64^, most motile cardiobots display high % of contractile areas up to 100% coverage. It is likely that non-optimized contractions, as seen in cardiobots, require larger areas of muscle tissue to generate comparable thrust. In evolutionary terms, this is in line with recent findings or hypotheses of early muscular-based motile animals, where it is reported that muscles were a high percentage of area in the early body plans^20^.

In comparison to the movement patterns of anthrobots and Xenobots, where the motility is powered by ciliated cells, the motility of cardiobots seems more erratic and less continuous. Towards increasing efficiency of motility in cardiobots, protocols of training/conditioning with paced contractions could be useful. Or, generation of body plans with more controlled contraction and attachment sites could also increase efficiency, as currently they are completely left as self-organized and not controlled. To sustain more stable motility, increased control of the body plan may be necessary. Positioning of the cardiobots, given that they are complex 3D objects, may affect the motility too.

The multivariate analysis provided some hints on what constitutes a strong body plan for motility. We show that high contraction area is highly correlated with high motility, as well as a body shape that can be either more or less spherical. One caveat with this analysis is the large variability of the dataset, since even for the same protocols, the variation of the outcome is rather high. We also noticed a trend, as in many other differentiation protocols, that experiment-to-experiment variation can be significant, both for differentiation markers, and for motility assays. This can be attributed to different batches of growth factors, or to hard-to-control parameters, like initial cell numbers. Especially for the extended protocol, the correlation analysis with the track length gave a non-null correlation with experiment id, signifying that certain experiments recorded higher motility scores compared to others. Importantly, synthetic organizer based structures seem to suffer from less experiment-to experiment variability. It will be important in future studies to optimize protocols to increase reproducibility within experiment and inter-experiment. Generating aggregates with more controlled morphological and patterning features would likely improve the consistency and the stability of the trajectories.

We also share that it may be possible that developmental trajectories like the cardiobots ones described here can shed some light on the evolution of development of motile animals^11,12,14–18^. What were the early developmental trajectories that generated the first body plans that showed muscle-based motility? Although it might turn out to be impossible to know exactly how they looked, it is tempting to speculate that they may have looked not dissimilar to the ones described here. Contractile cells likely existed before the first contractile multicellular animal^23^, so the co-option of their differentiation for multicellular body movement could have been supported by one of the first differentiations and division of labor between support cells and contractile cells.

In this scenario of course there are several differences between the ancestral animal and cardiobots. In cardiobots, for example, contraction is highly optimized in cardiomyocyte-like cells, whereas contractile cells in ancestral animals were probably far less optimized. Their localization then must have been quite precisely arranged to generate the thrust for multicellular motility, to out-compete cilia-based motility, which could reach very high speeds at small dimensions. The biggest advantage for muscle-based contractility is identified in capacity to move larger body masses.

In cardiobots, where cardiomyocytes are used for motility, we have in a sense inverted an hypothesized evolutionary transition: muscle-based locomotion is thought to predate formation of a circulatory system^65^. In evolutionary terms, it is interesting to speculate on the relationship between these two contractile systems. Genetic circuits that were initially utilized for generating patterned contractility for locomotory function, might have been repurposed to generate contraction in conjunction with tubulogenesis of the early blood vessels towards cardiovascular system development. In line with this, it seems that skeletal muscle and cardio-muscles evolved from common precursors^66,67^. It will be interesting to evaluate which innovation at the origin of the cardiovascular system enabled the repurposing of locomotory contractility developmental circuits for cardiovascular purposes. For example, enhancers of cardiogenic factors like Nkx2.5, GATA4, and Tbx5 have binding sites for signaling effectors such as Smads (from BMP signaling), MEF2, and bHLH factors (e.g., MyoD family), which are also crucial for skeletal muscle differentiation and patterning^68,69^. It remains to be explored if such regulatory cross-utilization represents a conserved evolutionary feature, as suggested by comparative studies in basal chordates and non-vertebrate deuterostomes. The fact that cardiobots display autonomous motility suggests that, to a certain extent, skeletal and cardiac contractility can be interchangeable.

In this study, we have shown an exploration, characterization and identification of self-organization protocols for the generation of muscle-based motile aggregates from stem cells. The combination of the developmental engineering design with quantitative benchmarks of physiological performance employed here is broadly applicable to reverse engineering muscular organs or simple life forms that display contractility-mediated motility with different architectures and capabilities. We hope these studies inspire and motivate the next generation of developmental engineering research.

## Acknowledgements

We thank all members of the Morsut Lab, Dr. Megan McCain, Dr. Matt Thomson, Dr. Mattia Gazzola for their valuable feedback and discussions throughout the course of this project. We also thank Sammana Kedir and Manasee Ramchandra Parulekar for writing the code used to perform the principal component analysis (PCA). We thank Inna Strazhnik for the illustration of Fig. 1B. We further thank Dr. Seth Ruffins and the Optical Imaging Facility for assistance with microscopy and Dr. Bernadette Masinsin, Dr. Jorge Contreras and the Flow Cytometry Facility for help with FACS experiments.

Research reported in this publication was supported by NIGMS of the National Institutes of Health under award number R35GM138256 (L.M.); the National Science Foundation award number CBET-2145528 Faculty Early Career Development Program (L.M.); NSF RECODE from CBET-2034495 (L.M.); USC Department of Stem Cell Biology and Regenerative Medicine Startup Fund (L.M.); Wellcome Trust, Leap HOPE (L.M.), Viterbi Center for CIEBOrg (L.M.). grant 2023-332386 from the Chan Zuckerberg Initiative Donor Advised Fund, CZI DAF, an advised fund of the Silicon Valley Community Foundation (LM). CIRM training grant EDUC4-12756 (F.G.). BAEF fellowship (BS). CIRM Bridges fellowship (KP).

## Author Contributions

CH and LM conceived the study, designed the experiments and wrote the manuscript. CH performed the experiments and analyzed the data. LM directed the research and acquired funding. F.G built the synthetic organizers, characterized them and contributed to the immunostaining experiments and its quantifications. GG contributed to protocol optimization and motility assays; morphological and contractility feature identification and quantification; Calcium experiments including quantifications; help with the initial setup of IF experiments. KP contributed to FACS, immunostaining, and Calcium experiments including quantifications. MK performed PCA analysis and correlation matrices. BS contributed to the FACS experiments and optimization. SM contributed to the motility assays and immunostaining.

## Videos

**Video 1-3**: Representative videomicroscopy of calcium transients in cardiobots at day 10 of differentiation, visualized using the calcium-sensitive dye Fluo-4. Timelapse duration: 34 seconds. Scale bar: 200 µm.

## Methods

### Material and methods

#### Cell lines

T-Brachyury eGFP mESC line was provided courtesy of Keller’s lab. The NKX2-5 emGFP mESC line was purchased from MMCA and generated as described here^53^. The dual reporter cell line, T-Brachyury_Sox17 Strawberry mESC line, was provided by Nachman’s lab and was generated as described here^49^.

#### Mouse embryonic stem cell maintenance

Mouse embryonic stem cells (mESCs) were cultured in 2i LIF medium consisting in DMEM (Thermo Scientific 11965112) supplemented with 10% of Fetal bovine serum (Cytiva SH30071.03), 1x Non essential amino acid (Thermo Fisher Scientific 11140050), 1X of Glutamax Supplement (Thermo Fisher Scientific 35050061), 0.1 mM β-Mercaptoethanol (Sigma-Aldrich M3148-25ML), and 1% Penicillin-Streptomycin (Thermo Fisher Scientific 15140122). The medium was further supplemented with 3 µM CHIR99021 (Stemcell Technologies 72054), 1 µM PD0325901 (Stemcell Technologies 72184), and 1000 U/mL LIF (Santa Cruz Biotechnology sc-4989A). Cells were plated on gelatin-coated dishes and maintained at 37°C with daily medium changes. Cells were passaged every 2-3 days using accutase (Thermo Fisher Scientific A1110501)* for 4min in 37°C before reaching 80-90% confluence.

#### Plasmid Construction

*pSpCas9-ROGI1* and *pHDR-ROGI1-MCS* were made in-house following published resources^71,72^. The *EF1a* promoter and *Cdh3-pA* were PCR-amplified from an pHDR-ROSA26-MCS and P-cells gDNA ^73^ respectively, and cloned into BamHI-digested *pHDR-ROGI1-MCS* using Gibson Assembly. *pSpCas9-ROSA26* and *pHDR-ROSA26-MCS* were a gift from Jamie Davies and have been described previously. *TagBFP-2A* was assembled with either *2A-Bmp4* or *2A-Wnt3a*, and *turboGFP-2A* was assembled with either *2A-VEGF165*, *2A-FGF2*, or *2A-FGF10*, together with KpnI-digested *pHDR-ROSA26-MCS* using Gibson Assembly. *TagBFP* was amplified from an pTH11 in-house construct; *turboGFP* was amplified from E-cells gDNA^73^; *Wnt3a* was amplified from HEKCdh3-Wnt3a gDNA^28^; *Bmp4, VEGF165, FGF2, FGF10* were outsourced as synthetic gene fragments (Twist Bioscience). All PCR amplifications were performed using 1-2ng input for plasmids, 100-200ng input for gDNA, and 2x Q5 DNA polymerase master mix in total reaction volumes of 25μl. All assemblies were performed using 30fmol vector, 50fmol per insert, and 2x HiFi DNA assembly mix in total reaction volumes of 20μl at 50C for 1 hour. Stellar competent bacteria were transformed with reaction volumes corresponding to 2fmol vector via heat shock method (30 minutes on ice, 30 seconds at 42C), 9 parts of SOC medium were added to 1 part bacteria, transformants were outgrown at 37C 250rpm for 40 minutes, and 1/10th of the outgrowth was plated on ampicillin-supplemented agar plates. Colony-derived plasmids were screened via restriction digestion followed by full plasmid sequencing (Primordium labs).

#### Synthetic Organizer Engineering

T-REx-293 cells were cultured in DMEM supplemented with 10% FBS, 2mM L-glutamine, 1% Penicillin/Streptomycin, and 0.1mM β-mercaptoethanol. Cells were harvested via regular trypsinization followed by neutralization using culture medium at 80-90% confluence, and passaged in a 1:10-1:12 ratio every 3-4 days. Cells were engineered to express *Cdh3* constitutively from the *ROGI1* locus by co-transfecting them with *pHDR-ROGI1-Cdh3* and *pSpCas9-ROGI1* using Lipofectamine3000. Transfections were set-up in 6-well plates containing T-REx-293 cells at 70-80% confluence. Five microliters Lipofectamine3000 reagent was added to 145μl OptiMEM and the solution was vortexed. Three micrograms of *pHDR-ROGI1-Cdh3* and 1μg *pSpCas9-ROGI1* was added to 145μl OptiMEM, and 5μl P3000 reagent was added. The DNA-P3000 solution was mixed into the Lipofectamine solution and incubated on site for 15 minutes. T-REx-293 cultures were aspirated of regular culture medium, gently washed in PBS, and refed with 1 ml OptiMEM. The transfection solution was distributed drop-wise across the well while rocking the plate and returned to the 37C, 5% CO2 incubator. After 6 hours of incubation, 2 ml of culture medium was added. Three days after transfection, culture medium was replenished with fresh medium containing G418 at 800μg/ml, and replenished every other day for 14 days. *Cdh3*-expressing T-REx-293 cells were subsequently engineered to express fluorescently-reported ligands (*Bmp4*, *Wnt3a*, *FGF2*, *FGF10*, *VEGF165*) from the *ROSA26* locus in a doxycycline-inducible manner. Transfections were performed as above using 3μg *pHDR-ROSA26-ligand* and 1μg *pSpCas9-ROSA26*; the medium was replenished with fresh medium containing 10μg/ml puromycin 3 days after transfection, and replenished every other day for 9-12 days.

#### Synthetic Organizer Characterization

Two-hundred-thousand cells were seeded in 12-well plates in culture medium containing doxycycline between 10pM - 100nM at 10-fold increments. Twenty-two hours after seeding, cells were harvested via standard trypsinization, pelleted, resuspended in 0.6ml PBS-2% FBS, and filtered through flow cytometry tube cap filters. Samples were analyzed on a BD Symphony cytometer for mCherry, TagBFP, and turboGFP fluorescence following gating of single cells, with non-engineered T-REx-293 and no-doxycycline synthetic organizer lines used as controls.

#### Synthetic Organizer-Stem cell co-cultures

Synthetic organizer cells were harvested, counted using 0.1% Trypan Blue in a Countess-II FL Automated Cell Counter, desired numbers were resuspended in serum-free N2B27 medium containing 500nM doxycycline, accounting for 500 cells per 50μl medium. Fifty microliters (500 cells) were seeded in low-adhesion U-shaped 96-wells per organizer. Bmp4 and Wnt3a organizers were prepared in parallel to stem cell seeding for differentiation (day 0), whereas VEGF, FGF2, FGF10 organizers were prepared 72h after stem cell seeding (day 3). Two days after organizer cell seeding, organizer spheroids in 50μl were manually transferred to stem cell aggregates in 50μl, and differentiations were continued as described herein with the addition of 500nM doxycycline in every medium change step.

#### Generation of cardiogenic gastruloid and novel cardiobots protocols

Standard gastruloids were generated using the protocol described by Van den Brink et al. (2014), while cardiogenic gastruloids were generated following the method described by Rossi et al. (2021). In both cases, samples were kept in 96-well plates at 37°C, 5°C in an incubator until 168h of differentiation, then transferred to an orbital shaker set to 40 rpm in 6-well plates. Briefly, 250 mESC were seeded in Nunclon Sphera 96 well U-bottom plates (Thermo Scientific 174929) in N2B27 medium. The differentiation medium is serum free and contained (1:1 neurobasal medium (Thermo Scientific 21103049) and DMEM/F12 (Thermo Fisher Scientific 11320033) supplemented with 1X B27 (50X) (Thermo Fisher Scientific 17504044) and1X N2 supplement (Thero Fisher scientific 17502-048), 100U/mL of penicillin-Streptomycin (Thermo Fisher Scientific 15140122), 2mM GlutaMAX (Thermo Fisher Scientific 35050061), 0.1mM MEM Non-essential Amino Acid (100X) (Thermo Fisher Scientific 11140050), 1mM Sodium Pyruvate (Thermo Scientific 11360070) and b-mercaptoethanol (Sigma M3148). Cells were then aggregated for 48h before adding a pulse of 3uM of CHIR99021(Stemcell Technologies 72054) for 24 hours. Afterward, Chiron is washed off and cells remain in N2B27.

For novel cardiogenic embryoid bodies, we replaced mesoderm induction with 3uM of CHIR99021 using a combination of three factors: 5ng/mL Activin A (Proteintech HZ-1138), 0,25ng/mL BMP-4 (Proteintech HZ-1045), and 5ng/mL VEGF (Fisher Scientific 293-VE-010/CF). After a pulse of these tri-factors (ABV), the medium was replaced with a factor-free N2B27 medium. For a second induction to enhance cardiac specification between 96 and 144 hours, the N2B27 basal medium was replaced with StemPro medium (Thermo Fisher Scientific 10639011) supplemented with 0.45nM L-Ascorbic acid (Sigma-Aldrich 255564-100G), 5ng/mL VEGF, 5ng/mL FGF2 (R&D Systems 233-FB-010/CF), and 25 ng/mL FGF10 (R&D Systems 345-FG-025/CF). This specific medium, termed cardiomyocyte differentiation medium (CDM), has been published here.

#### Aggregates imaging and quantifications

To monitor the differentiation everyday. U Shaped bottom 96 well plate have been scanned using AxioZeiss Observer. Images have been collected and pre-processing using Fiji/ImageJ. Area, eccentricity, length of the Embryos bodies have been segmented and quantified using MOrgAna pipeline (described 34494114).

#### Calcium Imaging and analysis and contractile dynamics

To monitor calcium dynamics, embryoid bodies were incubated with 0.25 ng/mL Fluo-4, AM (Thermo Fisher Scientific, F14201) in culture medium for 40 minutes at 37°C on an orbital shaker. Following incubation, the dye-containing medium was replaced with fresh medium to remove unincorporated probe prior to imaging. Time-lapse imaging was performed using a Zeiss Axio Observer equipped with a Stream camera and GFP filter settings to detect calcium-dependent fluorescence.

High-frequency 1-minute time-lapse sequences were acquired to capture rhythmic fluorescence changes corresponding to contractile events. Fiji (ImageJ) was used to process the fluorescence data, and oscillatory signals were extracted and analyzed using wavelet transformation as previously described^74^.

Regions of periodic calcium activity were quantified to determine the area of contraction relative to total aggregate area. Displacement per contraction was calculated by tracking tissue movement during individual contraction cycles. Calcium imaging thus served to delineate the spatial extent of contraction and quantify contraction eccentricity, which was subsequently included in principal component analysis (PCA) for phenotypic clustering.

#### Motility assays and analysis

After 7 days of differentiation, samples were transferred from a 96-well plate into a low-adherence plate, with approximately 6 to 10 samples per well of a 6-well low-adherence plate (Corning 3471) for imaging.

Time-lapse imaging of the samples was performed using a Keyence microscope (BZ-X810). One picture was taken every 5 minutes for at least 30 minutes using 1X magnification (Nikon objective), high resolution, and brightfield. Samples at day 10 followed a similar procedure, except they were left on the orbital shaker at 40 rpm for 2 days. For data analysis, Imaris software was used to track each aggregate and quantify them over time.

#### Immunostaining and quantification

Embryoid bodies were collected at 120 hours (Day 5), 168 hours (Day 7), and/or 240 hours (Day 10) of differentiation. Aggregates were washed multiple times with phosphate-buffered saline (PBS) to remove residual culture medium, then fixed in 4% paraformaldehyde (PFA) for 1 hour at room temperature. After fixation, samples were permeabilized twice for 30 minutes on ice in 0.5% Triton X-100 while gently rocking. Blocking was performed using a solution of 1% bovine serum albumin (BSA) and 1% Triton X-100 for 1–2 hours on ice. Aggregates were subsequently incubated with primary antibodies overnight at 4°C on a nutating mixer. The primary antibodies used included: anti-cardiac Troponin T (1:400, Thermo Fisher Scientific, MA5-12960), anti-SOX2 (1:400, Abcam ab79351), anti-EOMES (1:200, Abcam ab23345), and anti-SOX17 (1:400, R&D Systems AF1924). Following three 30-minute washes with PBS on ice, samples were incubated overnight with Alexa Fluor 647-conjugated goat secondary antibodies (1:500, Invitrogen A21236) and DAPI for nuclear staining. Final washes (3 × 30 minutes) were performed on ice before transferring aggregates to 6-well plates for imaging. Images were acquired using a Zeiss Axio Observer microscope or Leica SP8. Image processing and quantification were performed using Fiji (ImageJ). Cardiac Troponin T (cTnT) fluorescence intensity was calculated as the ratio of total cTnT signal intensity to the total DAPI-positive area of the aggregate.

#### Flow cytometry and analysis

For time course experiments to quantify different germ layers, embryoid aggregates were collected at specified time points of differentiation, and washed with dPBS. Samples were incubated with 3 mM EDTA in dPBS at 37°C for 5-7 mins, then mechanically dissociated. Cells were subsequently neutralized with DMEM and washed with 1% BSA in dPBS. Nkx2-5, Sox17 and T-brachyury expression were measured with reporter fluorescence levels (eGFP and mCherry) at the Attune at the USC flow cytometry core.

To quantify cardiac progenitors Flk1/PDGFRa, T Brachyury-eGFP embryoid bodies were collected and dissociated at 96h and 120h of differentiation as described above. Then, samples were first incubated with a Flk1-biotinylated antibody (Invitrogen 13-5821-82) (dilution 1:100) for 30 mins in FACS buffer (1% BSA in PBS) on ice. Samples were then rinsed to remove the remaining antibodies. After this rinsing step, this was followed by another 30 mins of incubation on ice with an antibody cocktail consisting of streptavidin-phosphatidylethanolamine PE-Cy7 (BD Pharmingen 557598) (Dilution 1:400) and a PDGFRa PE-conjugated antibody (Invitrogen 12-1401-81) (Dilution 1:75) in FACS buffer. Stained cells were subsequently washed as previously described above. All cells were resuspended in FACS buffer, 1% BSA in dPBS with DAPI (1:10000), then measured with the ThermoFisher Attune NxT Flow Cytometer. FACS data was analyzed using FlowJo.

##### Tracker measurements

Contraction length and displacement per contraction were measured using the free, open-source physics video analysis software Tracker (https://opensourcephysics.github.io/tracker-online/). Captured videos were first converted into .avi files and set to 19.52 frames/sec. Prior to analysis, video scaling was achieved using calibration sticks set to 2.498e-3 m at −180.0° and 2.005e-3 m at −270.7°. Coordinate axes were then set to the target location (contracting pole or non-contracting pole) on the aggregate. The point mass feature was used to autotrack the position of the target location for the duration of the video. Coordinates were converted to distances using the distance formula. Contraction length was calculated by measuring the average length difference between relaxation and contraction throughout a recording. Displacement per contraction was measured from a non-contracting pole, with its final distance normalized by the number of times the aggregate contracted within the video.

#### Data Analysis

To investigate the multidimensional relationships between morphology, contractility, and motility phenotypes in embryoid aggregates, we performed both correlation matrix analysis and principal component analysis (PCA) with unsupervised clustering.

#### Correlation Matrix

A pairwise Pearson correlation matrix was constructed using Python (v3.11.5) with pandas, numpy, matplotlib, and seaborn. The dataset included contractile and morphological features extracted from the above imaging and motility assays. Preprocessing steps included 1. Removal of non-contracting aggregates (Contracting_Area_Percent = 0). 2. Exclusion of samples from ExpID = 12 and Protocol = 2 (Chiron-based). 3. Conversion of relevant columns to numeric values with NA coercion. 4. Removal of auxiliary metadata (ExpID, Protocol, and derived metrics such as Track_Straightness and Avg_Speed). The resulting dataset was cleaned and passed into a correlation matrix function that used df.corr() to compute Pearson coefficients. A heatmap of the correlation matrix was visualized using seaborn’s heatmap function with the coolwarm color scale, annotating all pairwise correlations and highlighting both positive and negative associations across contractility, morphology, and motility metrics. The figure was saved in scalable vector graphics (SVG) format for high-resolution publication.

#### Principal Component Analysis (PCA) and Clustering

To reduce dimensionality and identify phenotypic clusters, PCA was performed in R (v4.3.2) within RStudio (v2025.05.0+496) using the prcomp() function on z-score normalized data. The input dataset included 10 quantitative features spanning area, shape (aspect ratio, eccentricity, circularity), and contractility (frequency, contraction area, percent contracting area, eccentricity of contraction, and contractile length average). Multiple PCAs were conducted on different subsets of the data, including sets with only contractile aggregates, sets with and without motility features and sets with all features.

Dimensionality reduction via PCA was followed by k-means clustering, with the number of clusters (k = 3) determined by both the elbow and silhouette methods (fviz_nbclust() from factoextra). Each aggregate was assigned to a cluster (A, B, or C), and visualization of the clusters was done in PC1–PC2 space.

Cluster-level phenotypic characterization was performed using a range of visualizations generated in R with ggplot2, ggpubr, and supporting packages. These included, but were not limited to, violin plots with overlaid jittered scatter points to compare morphological, contractility, and motility features across PCA-defined clusters. Statistical comparisons were conducted using stat_compare_means() from the ggpubr package. Pseudo-logarithmic scaling was applied to variables with skewed distributions, such as total track length and displacement. Feature distributions were also visualized by protocol to identify trends associated with different experimental conditions. Principal component loadings were examined to determine which features contributed most to PC1 and PC2. Stacked bar plots were used to show how different protocols were represented across clusters and to assess protocol-associated shifts in phenotypic outcomes.

All plots were exported as SVG files via ggsave() to ensure compatibility with downstream figure assembly and publication formatting.

#### Data Analysis and Visualization

All data analyses and visualizations were performed using a combination of open-source and commercial software tools. FIJI (ImageJ) was used for image processing and quantitative analysis of immunofluorescence and calcium imaging data. Imaris (Bitplane) was utilized to analyze time-lapse video recordings, enabling the extraction of quantitative motility parameters, including track length, displacement length, and speed of embryoid body movement. Bar graphs and statistical visualizations were generated using GraphPad Prism 10.4.2 (534) Serial number: GraphPad Prism 10.4.2 (534). Expiration: Feb 3, 2026. Serial number:

GPS-2333022-L###-#####. Activation code:

ACTGP-54E4D140-F4FD7C85-817C5E85-F3300393. MachineID: 5C8EE2083D1

