## Supplemental figures and figure legends for "Derivation of cardiomyocyte-propelled motile aggregates from stem cells"

### SUPPLEMENTARY FIGURES

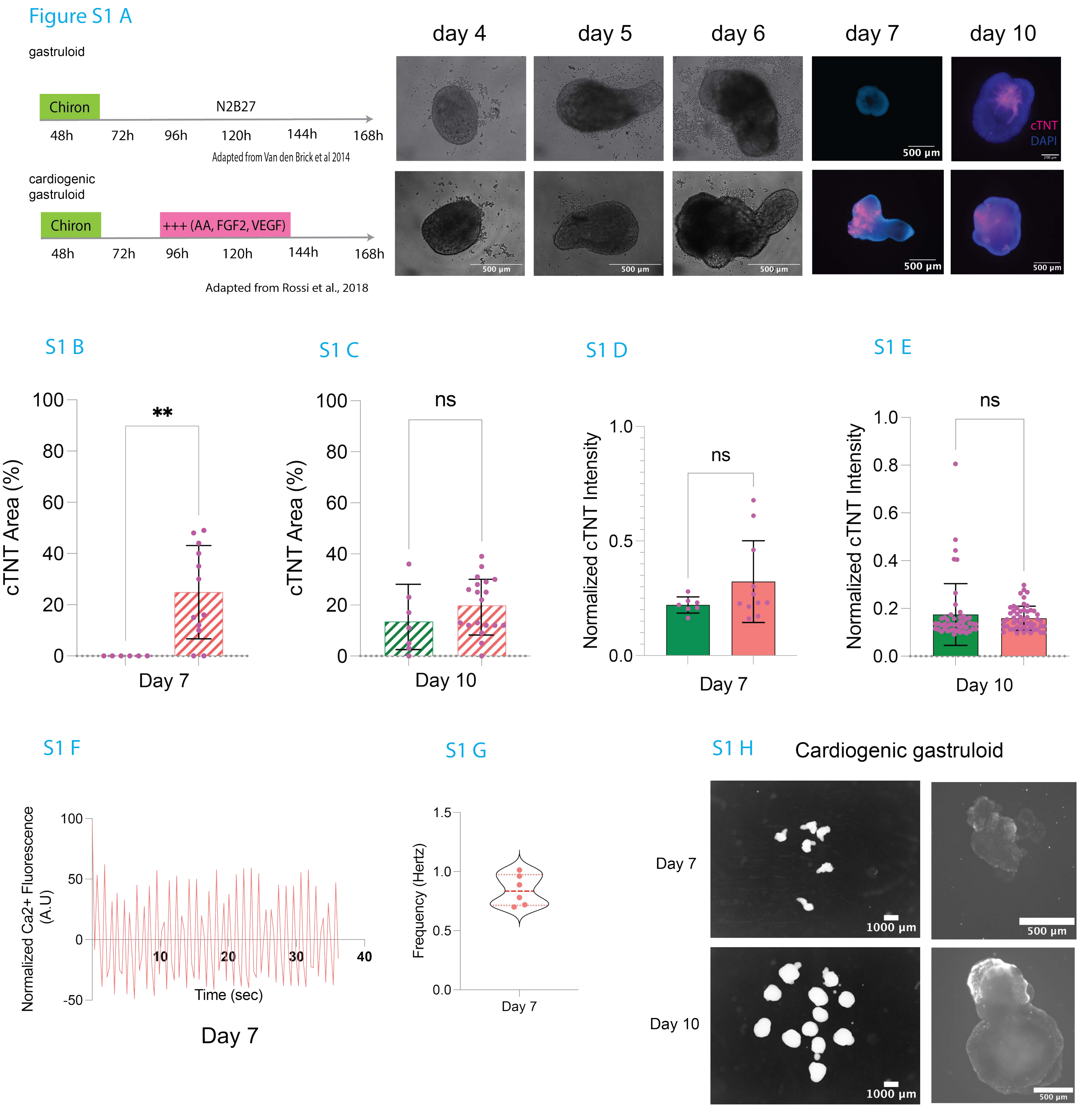


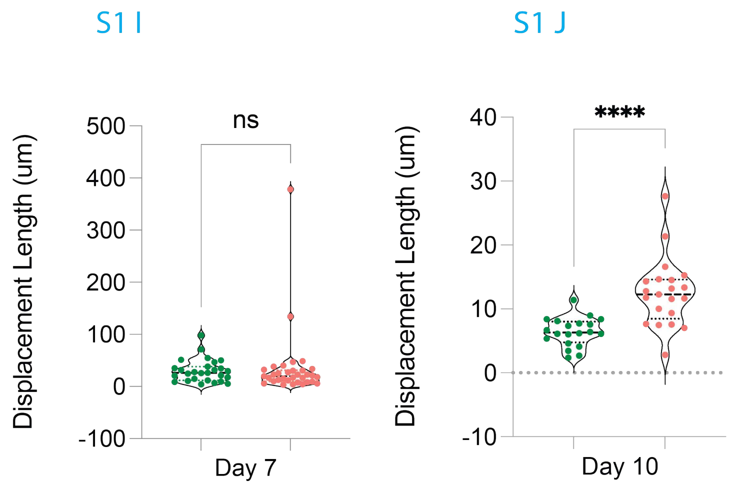


## S1 H

| Locomotion Type | Speed Range | Typical Range per Second |
|---|---|---|
| Ciliary swimming | 200–1000 µm/s | 0.5–2 body lengths/s |
| Flagellar swimming | 10–100 µm/s | 0.05–0.2 body lengths/s |
| Undulatory (e.g. worms) | 100–500 µm/s | 0.3–1.5 body lengths/s |
| Slithering / Surface gliding | 1–20 µm/s | very local (< 0.05 BL/s) |

### Figure Supplemental 1

**A.** Left: Schematic illustration of gastruloid and cardiogenic gastruloid protocols adapted from van den Brink et al., 2014, and Rossi et al., 2018. Right: day 4-6, microscope images of embryoid bodies at different time points captured using brightfield imaging. Day 7 and 10, fluorescent microscope images of aggregates immunostained with DAPI (blue) and cardiac troponin T (cTnT, magenta). Scale bar 500um. **B.** Bar graph showing the percentage of cTnT-positive area over the total area of individual aggregates on Day 7 for gastruloids (n=6) and cardiogenic gastruloids (n=12). *n* = number of individual aggregates. Statistical analysis was performed using the Mann–Whitney test. **C.** Bar graph showing the percentage of cTnT-positive area on Day 10 for gastruloids (n=7) and cardiogenic gastruloids (n=19). *n* = number of individual aggregates Statistical analysis was performed using the Mann–Whitney test. **D.** Bar graph showing normalized cTnT expression intensity on Day 7 for gastruloids (n=7) and cardiogenic gastruloids (n=11). *n* = number of individual aggregates. Statistical analysis was performed using the Mann–Whitney test. **E.** Bar graph showing normalized cTnT expression intensity on Day 10 for gastruloids (n=48) and cardiogenic gastruloids (n=52). *n* = number of individual aggregates. Statistical analysis was performed using the Mann–Whitney test. **F.** Representative calcium oscillation signal recorded in a contracting cardiogenic gastruloid at Day 7 using the Fluo-4 calcium indicator. See methods for details of live imaging signal processing. **G.** Violin plot showing contraction frequency (in Hertz) on Day 7 for cardiogenic gastruloids (n=6). Dotted red line represents the median. *n* = number of individual aggregates. **H.** left: Micrograph picture of a group of aggregates at the indicated time point, as setup for motility assay (scale bar 1000um). Right: individual aggregate microscope picture of aggregate after incubation with Fluo-4 calcium probe in cardiogenic gastruloids (scale bar 500um). **I.** Bar graph showing displacement length (in micrometers) on Day 7 measured in motility assays for gastruloids (n=27) and cardiogenic gastruloids (n=33). *n* = number of individual aggregates. Statistical analysis was performed using the Mann–Whitney test. **J.** Bar graph showing displacement length (in micrometers) on Day 10 measured in motility assays for gastruloids (n=20) and cardiogenic gastruloids (n=21). *n* = number of individual aggregates. Statistical analysis was performed using Welch's test. **K.** Summary table showing, for each locomotion type, the characteristic speed range and typical displacement values per second.
Statistical significance: *$p < 0.05$; **$p < 0.01$; ***$p < 0.001$.

### Figure S2 A

| Protocol # | Protocols | Induction (Day 2-3) | Induction (Day 4-6) | % Contracting aggregate (Day 7) |
|---|---|---|---|---|
| | Gastruloid | Chiron | N2B27 | 2.8 |
| #1 | Cardiogenic gastruloid | Chiron | N2B27 (AA, FGF2, FGF10, VEGF) | 25.68 |
| #2 | Modified cardiogenic gastruloid | Chiron | StemPro (AA, FGF2, FGF10, VEGF) | 19.50 |
| #3 # | AB_N2B27 | Activin A, BMP-4 | N2B27 | 10.80 |
| #4 | AB_CDM | Activin A, BMP-4 | StemPro (AA, FGF2, FGF10, VEGF) | 52 |
| #5 | ABV_N2B27 | Activin A, BMP-4, VEGF | N2B27 | 0 |
| #6 | Cardiogenic mesoendoderm | Activin A, BMP-4, VEGF | StemPro (AA, FGF2, FGF10, VEGF) | 75.07 |
| #7 | Mesoendoderm | Activin A, BMP-4, VEGF | StemPro (AA, No factors) | 23.88 |

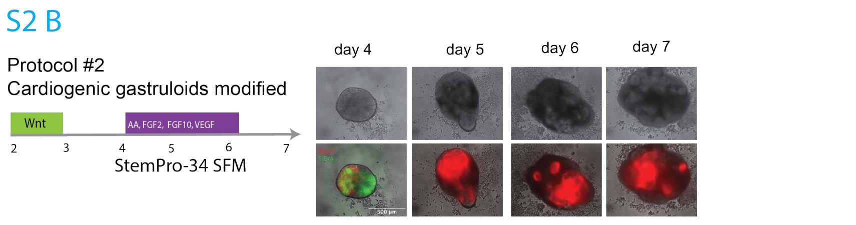


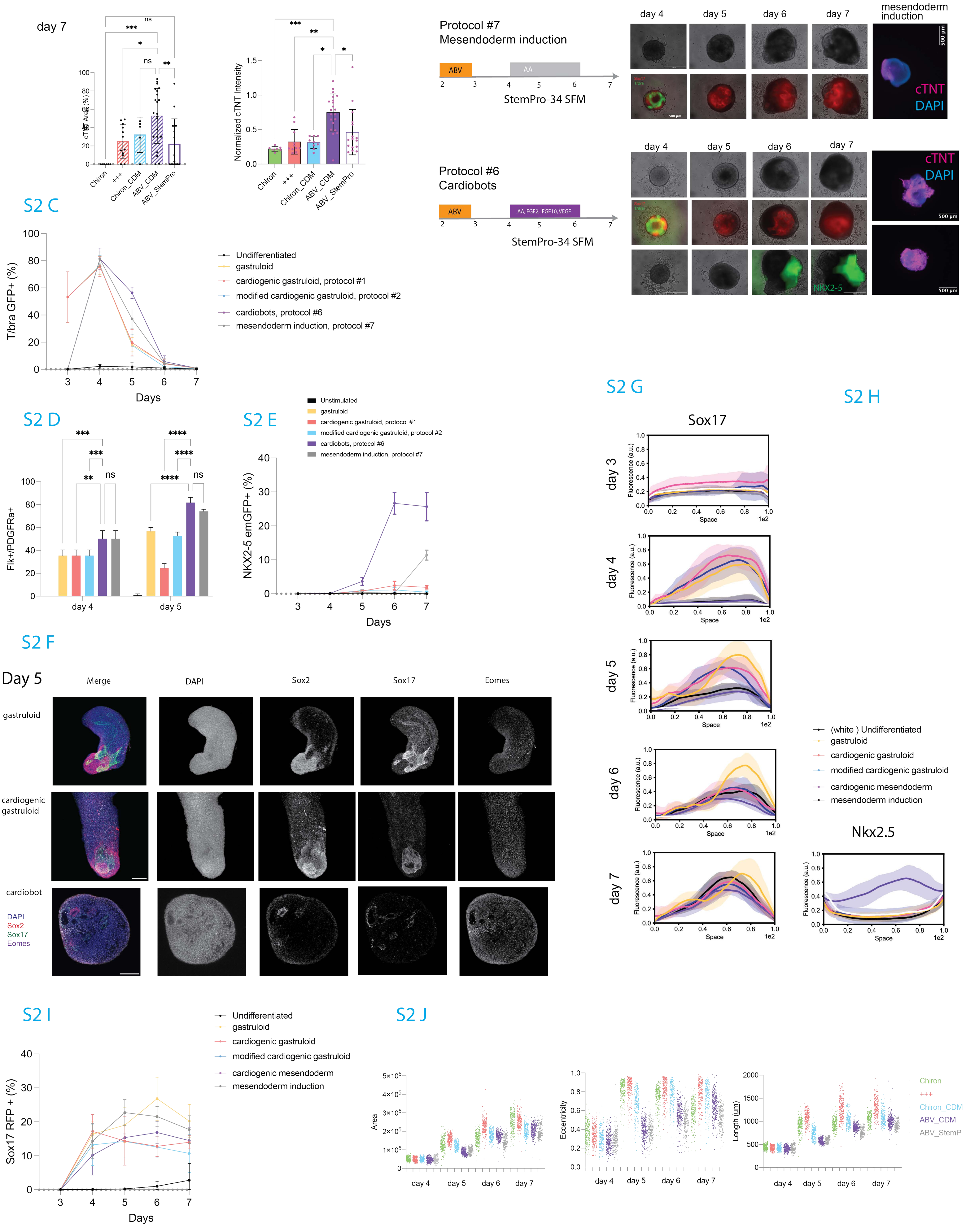

### Figure S2: Characterization and developmental trajectories of alternative cardiogenic embryoid body protocols.

**A.** *Top panel:* Summary table of tested cardiogenic protocols, indicating protocol titles, timing of induction, specific signaling factors used, and the percentage of contracting embryoid bodies observed by Day 7.
*Bottom panel:* Bar graphs showing the percentage of area positive for cardiac Troponin T (cTNT) and normalized cTNT fluorescence intensity for each protocol at Day 7. See methods for quantification procedures. Statistical analysis was performed using the Kruskal–Wallis test.

**B.** Left side: protocol schematics with the timeline and the factors used for mesoderm induction (days 2-3) and cardiogenic induction (days 4-6). Right side: Representative time-lapse images from Day 4 to Day 7 for protocols #2, #6, and #7. Images include brightfield views and fluorescence showing Sox17 (mStrawberry), T/Brachyury (eGFP), and NKX2-5 (emGFP, green).

**C.** Time course of average percentage of cells that express the T/Bra-GFP reporter over the total cells of individual aggregates of the indicated protocols.  Each dot is an average of N=5 independent experiments, where a minimum n=16 aggregates/per experiment and conditions have been used. Statistic tests have been performed using multiple t-tests.

**D.** Bar graph showing the average percentage of cells co-expressing FLK1 and PDGFRA at Day 4 and Day 5 of differentiation for each protocol. Each dot is an average of N=5 independent experiments, where a minimum n=16 aggregates/per experiment and conditions have been used. Statistic tests have been performed using multiple t-tests.

**E.** Time course of average percentage of cells that express the NKX2-5-GFP reporter over the total cells of individual aggregates, compared to unstimulated controls.  Each dot is an average of N=5 independent experiments, where a minimum n=16 aggregates/per experiment and conditions have been used. Statistic tests have been performed using multiple t-tests.

**F.** Related to Main Fig. 2H. Immunofluorescence images with overlay and single channel view of representative cardiogenic gastruloid, cardiobot and gastruloid samples at Day 5 stained for markers of germ layers: Sox2 (neuroectoderm, red), Sox17 (endoderm, green), EOMES (pan-mesendoderm, grey), and DAPI (blue). Scale bar: 100 µm.

**G-H.** Average Sox17-mCherry and Nkx2.5 reporter expression along the length of individual aggregates, for the indicated protocols in legend, at the indicated days.

**I.** FACS line plots showing the percentage of Sox17-RFP–positive cells over time in cardiogenic embryoid bodies versus unstimulated controls.

**J.** Dot plots showing distribution of morphological features—area, eccentricity, and length—across >200 individual embryoid bodies per protocol from Day 4 to Day 7.

Statistical significance: $*p < 0.05$; $**p < 0.01$; $***p < 0.001$.

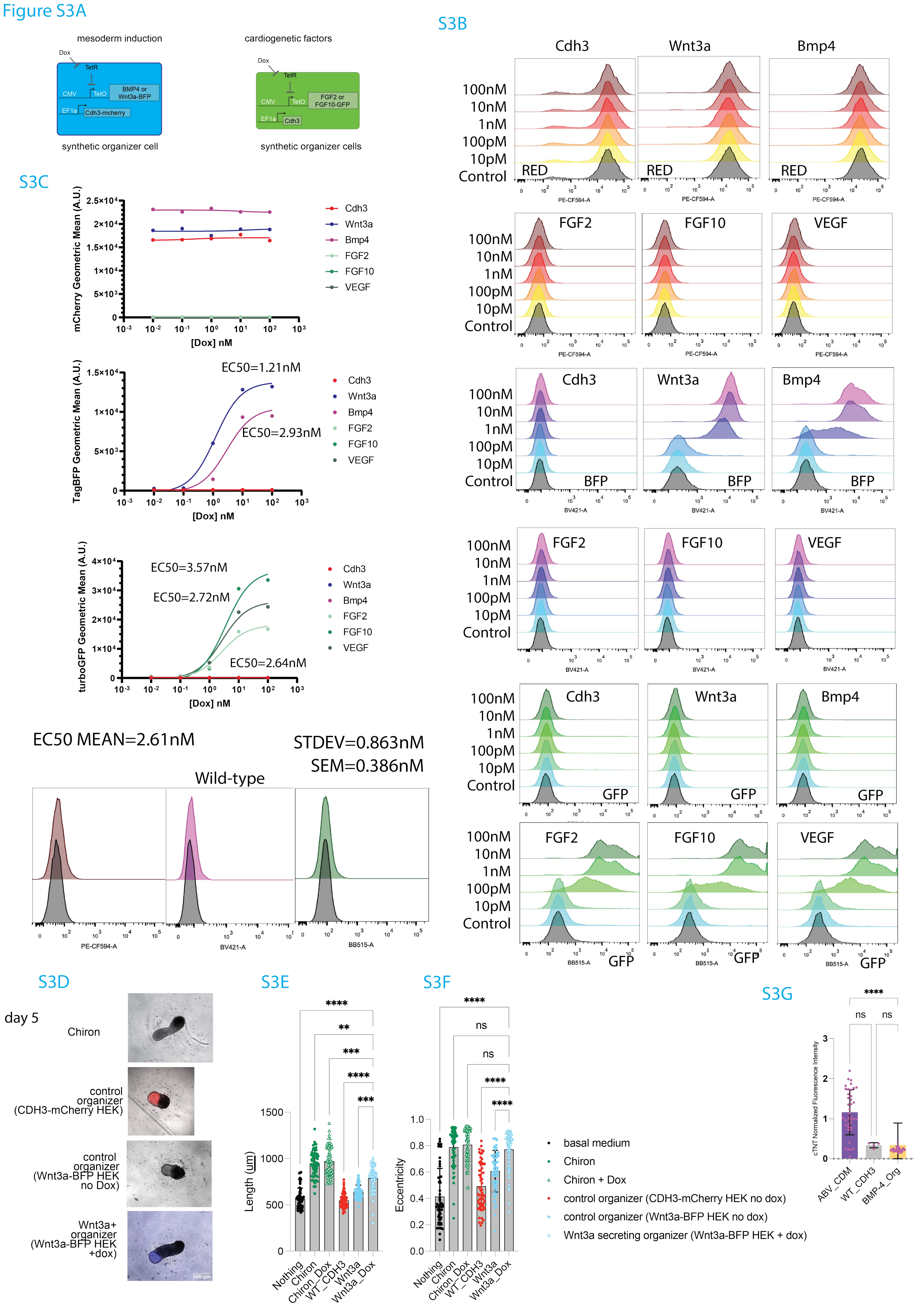
Figure S3A
mesoderm induction
cardiogenetic factors
Dox
TetR
CMV
TetO
BMP4 or Wnt3a-BFP
EF1a
Cdh3-mcherry
FGF2 or FGF10-GFP
Cdh3
synthetic organizer cell
synthetic organizer cells
S3B
Cdh3
Wnt3a
Bmp4
100nM
10nM
1nM
100pM
10pM
Control
RED
PE-CF594-A
FGF2
FGF10
VEGF
BFP
BV421-A
GFP
BB515-A
S3C
mCherry Geometric Mean (A.U.)
[Dox] nM
Cdh3
Wnt3a
Bmp4
FGF2
FGF10
VEGF
TagBFP Geometric Mean (A.U.)
EC50=1.21nM
EC50=2.93nM
turboGFP Geometric Mean (A.U.)
EC50=3.57nM
EC50=2.72nM
EC50=2.64nM
EC50 MEAN=2.61nM
STDEV=0.863nM
SEM=0.386nM
Wild-type
S3D
day 5
Chiron
control organizer (CDH3-mCherry HEK)
control organizer (Wnt3a-BFP HEK no Dox)
Wnt3a+ organizer (Wnt3a-BFP HEK +dox)
500 µm
S3E
Length (um)
Nothing
Chiron
Chiron_Dox
WT_CDH3
Wnt3a
Wnt3a_Dox
S3F
Eccentricity
ns
basal medium
Chiron
Chiron + Dox
control organizer (CDH3-mCherry HEK no dox)
control organizer (Wnt3a-BFP HEK no dox)
Wnt3a secreting organizer (Wnt3a-BFP HEK + dox)
S3G
cTNT Normalized Fluorescence Intensity
ABV_CDM
WT_CDH3
BMP-4_Org

### Figure S3: Characterisation of dox inducible synthetic organizers

**A**. Schematic illustrating the genetic constructs used to generate plasmids expressing either Wnt3a or BMP4, each linked to TagBFP and co-expressed with mCherry-Cdh3. Additional plasmids express turbo GFP-2A in association with either FGF2, FGF10, or VEGF, all co-expressed with Cdh3.

**B**. Fluorescence histograms showing expression levels of Cdh3, Wnt3a, BMP4, VEGF, FGF2, and FGF10 in organizer cells after 24 hours of exposure to the indicated doxycycline concentrations.

**C**. Dose-response curves showing the geometric mean fluorescence intensity (arbitrary units, AU) of mCherry, TagBFP, and turbGFP for different organizer constructs: Cdh3, Wnt3a, BMP4, FGF2, FGF10, and VEGF.

**D**. Representative snapshots of embryoid bodies at Day 5 of differentiation under various conditions: gastruloid, control organizer (Cdh3-mCherry), and assembloids containing Wnt3a organizer cells with or without doxycycline treatment.

**E**. Bar graph showing the length of embryoid bodies at Day 5 for the following conditions: basal medium only (black, $n$ = 57), Chiron without doxycycline (green, $n$ = 62), Chiron with doxycycline (green triangle, $n$ = 62), control organizer (red, $n$ = 59), Wnt3a organizer without doxycycline (blue, $n$ = 57), and Wnt3a organizer with doxycycline (blue triangle, $n$ = 55). $n$ = number of individual aggregates. Statistical analysis was performed using the Kruskal–Wallis test.

**F**. Bar graph showing the eccentricity of embryoid bodies at Day 5 for the same conditions listed in panel E. $n$ = number of individual aggregates. Statistical analysis was performed using the Kruskal–Wallis test.

**G**. Bar graph showing normalized cTnT expression intensity on Day 7 for the following conditions: cardiobots (purple, $n$ = 39), control organizer (grey, $n$ = 4), and BMP4 organizer (yellow, $n$ = 26). $n$ = number of individual aggregates. Statistical analysis was performed using the Kruskal–Wallis test.

Statistical significance: *$p < 0.05$; **$p < 0.01$; ***$p < 0.001$.

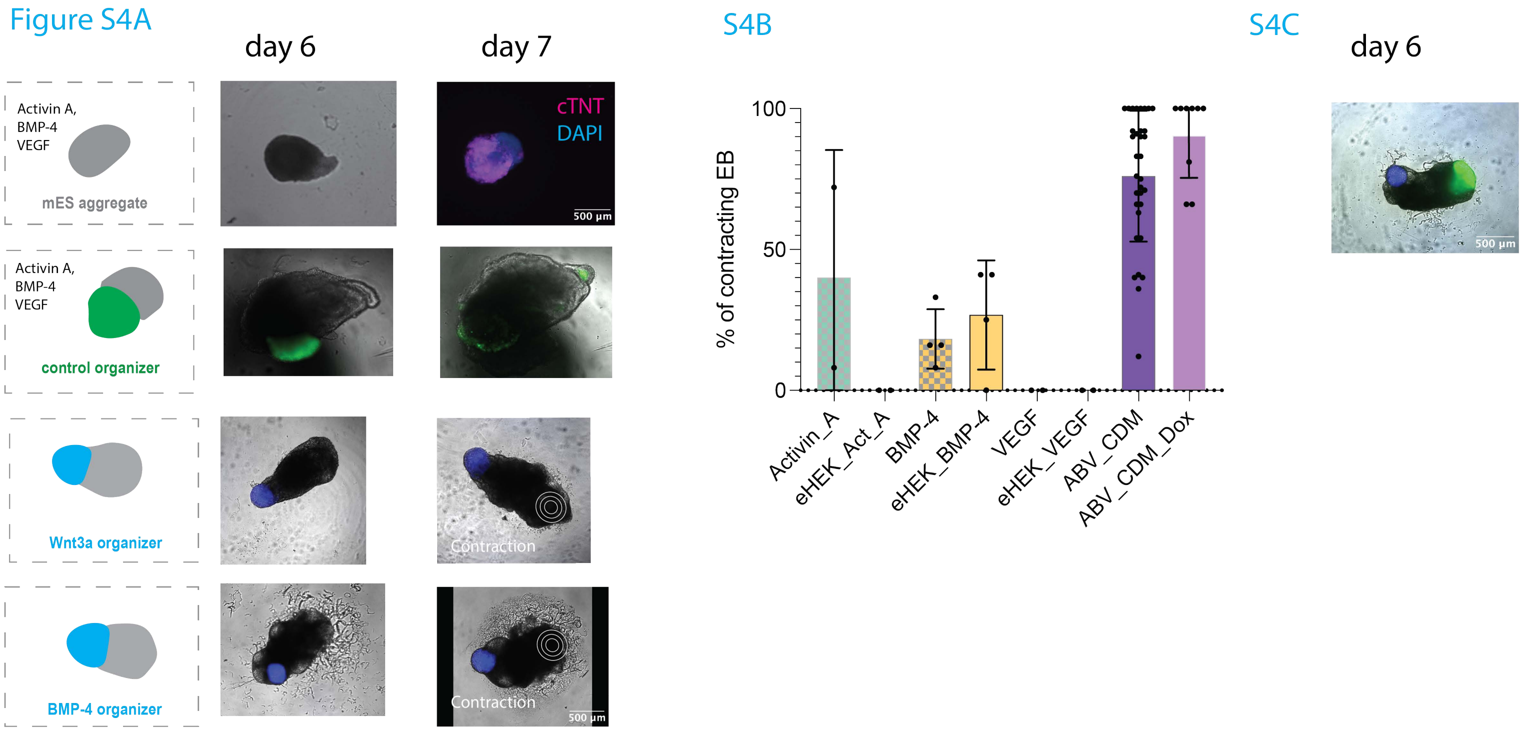
Figure S4A
day 6
day 7
Activin A,
BMP-4
VEGF
mES aggregate
cTNT
DAPI
500 µm
control organizer
Wnt3a organizer
Contraction
BMP-4 organizer
S4B
% of contracting EB
100
50
0
Activin_A
eHEK_Act_A
BMP-4
eHEK_BMP-4
VEGF
eHEK_VEGF
ABV_CDM
ABV_CDM_Dox
S4C
day 6

### Figure S4:

**A.** Representative immunofluorescence images of aggregates at Day 7, stained for cardiac Troponin T (cTNT, magenta) and nuclei (DAPI, blue). Scale bar: 500 µm. Representative snapshots of aggregates associated with Wnt3a organizers (blue, BFP reporter) or CDH3 (green). Concentric circles indicate the contracting regions.

**B.** Average percentage of contracting embryoid bodies in response to soluble factors—Activin A ($N = 2$), BMP4 ($N = 4$), VEGF ($N = 2$)—compared to their corresponding synthetic organizers (eHEK-Activin A, eHEK-BMP4, eHEK-VEGF), with ABV_CDM ($N = 6$) included as a control. *N* refers to the number of independent experiments.

**C.** Related to Main Fig. 3I, same conditions, datapoint for day 6.

Statistical significance: $^{*}p < 0.05$; $^{**}p < 0.01$; $^{***}p < 0.001$.

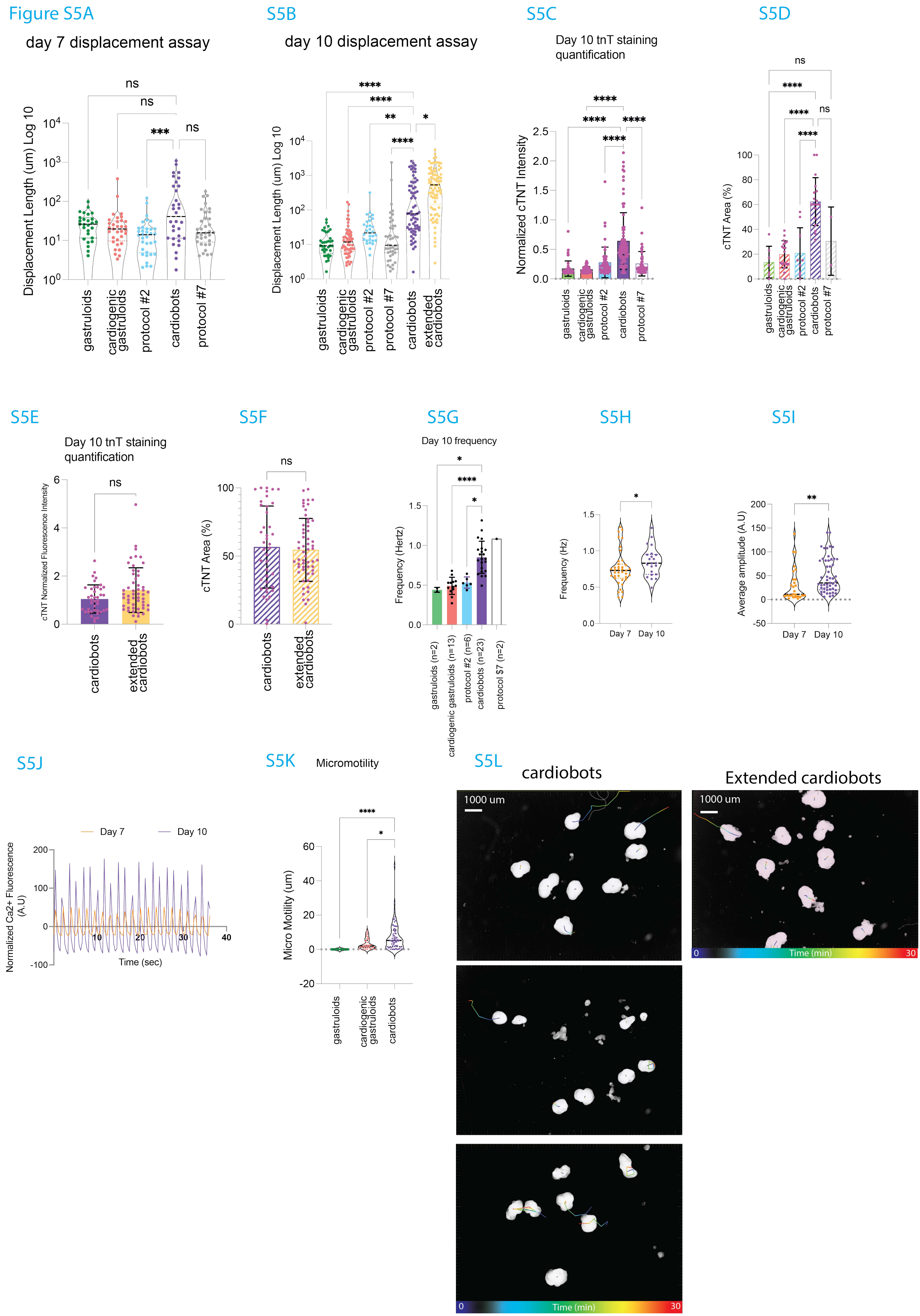

Figure S5A
day 7 displacement assay
Displacement Length (um) Log 10
gastruloids
cardiogenic gastruloids
protocol #2
cardiobots
protocol #7
ns
***
S5B
day 10 displacement assay
Displacement Length (um) Log 10
gastruloids
cardiogenic gastruloids
protocol #2
protocol #7
cardiobots
extended cardiobots
****
**
*
S5C
Day 10 tnT staining quantification
Normalized cTNT Intensity
gastruloids
cardiogenic gastruloids
protocol #2
cardiobots
protocol #7
S5D
cTNT Area (%)
gastruloids
cardiogenic gastruloids
protocol #2
cardiobots
protocol #7
S5E
Day 10 tnT staining quantification
cTNT Normalized Fluorescence Intensity
cardiobots
extended cardiobots
S5F
cTNT Area (%)
cardiobots
extended cardiobots
S5G
Day 10 frequency
Frequency (Hertz)
gastruloids (n=2)
cardiogenic gastruloids (n=13)
protocol #2 (n=6)
cardiobots (n=23)
protocol $7 (n=2)
S5H
Frequency (Hz)
Day 7
Day 10
S5I
Average amplitude (A.U)
Day 7
Day 10
S5J
Day 7
Day 10
Normalized Ca2+ Fluorescence (A.U)
Time (sec)
S5K
Micromotility
Micro Motility (um)
gastruloids
cardiogenic gastruloids
cardiobots
S5L
cardiobots
Extended cardiobots
1000 um
Time (min)
0
30

### Figure S5: Further Characterisation of motile aggregates: cardiobots

**A.** Violin plots of displacement length (µm, $log_{10}$ scale) of embryoid bodies at Day 7, measured over 30-minute motility assays under the following conditions: gastruloids (green, n = 17), cardiogenic gastruloids (red, n = 33), Chiron_CDM, protocol #2 (blue, n = 34), cardiobots (dark green, n = 32), and protocol #7 (grey, n = 34). Statistical analysis was performed using the Kruskal–Wallis test.

**B.** Violin plots of displacement length (µm, $log_{10}$ scale) of embryoid bodies at Day 10, measured over 30-minute motility assays under the following conditions: gastruloids (green, n = 37), cardiogenic gastruloids (red, n = 55), Chiron_CDM, protocol #2 (blue, n = 31), cardiobots (dark green, n = 68), and protocol #7 (grey, n = 44), extended cardiobots (yellow, *n* = 81). Statistical analysis was performed using the Kruskal–Wallis test.

**C.** Normalized cTNT fluorescence intensity in aggregates at Day 10. Conditions: gastruloids (*n* = 48), cardiogenic gastruloids (*n* = 52), Chiron_CDM, protocol #2 (*n* = 53), cardiobots (*n* = 80), and ABV_StemPro, protocol #7 (*n* = 46). *n* refers to the number of individual aggregates. Statistical analysis was performed using the Kruskal–Wallis test.

**D.** Average percentage of cTNT-positive area in aggregates at Day 10. Conditions: gastruloids (*n* = 7), cardiogenic gastruloids (*n* = 19), Chiron_CDM, protocol #2 (*n* = 9), cardiobots (*n* = 18), and ABV_StemPro, protocol #7 (*n* = 2). Statistical analysis was performed using one-way ANOVA.

**E.** Normalized cTNT fluorescence intensity in embryoid bodies at Day 10. Conditions: cardiobots (*n* = 40) and extended cardiobots (*n* = 53). Statistical analysis was performed using the Mann–Whitney test.

**F.** Percentage of cTNT-positive area in embryoid bodies at Day 10. Conditions: cardiobots (*n* = 40) and extended cardiobots (*n* = 57). Statistical analysis was performed using an unpaired *t*-test.

**G.** Calcium oscillation frequency (Hz) in contractile embryoid bodies at day 10 generated with the following protocols: gastruloids (green, *n* = 2), cardiogenic gastruloids (red, *n* = 13), Chiron_CDM, protocol #2 (blue, *n* = 6), cardiobots (purple, *n* = 23), and ABV_StemPro, protocol #7 (transparent, *n* = 2). Statistical analysis was performed using the Kruskal–Wallis test.

**H.** Violin plots comparing contraction frequencies (Hz) of contractile cardiobots at Day 7 (orange, *n* = 39) and Day 10 (purple, *n* = 22). *n* refers to the number of individual aggregates. Statistical analysis was performed using the Mann–Whitney test.

**I.** Average amplitude of calcium oscillations measured in contracting cardiobots at Day 7 (orange, *n* = 27) and Day 10 (purple, *n* = 53). Statistical analysis was performed using the Mann–Whitney test.

**J.** Representative normalized calcium oscillation traces recorded in contracting cardiobots at Day 7 (orange line) and Day 10 (purple line), using the Fluo-4 calcium indicator.

**K.** Violin plot of length of motility tracks during 30'' observations for aggregates generated with the indicated protocols.

**L.** Snapshots from motility assays for 30' recordings, done at Day 10 for the indicated protocols. Scale bar: 1000 µm.

Statistical significance: *$p < 0.05$; **$p < 0.01$; ***$p < 0.001$.

#### Figure S6A

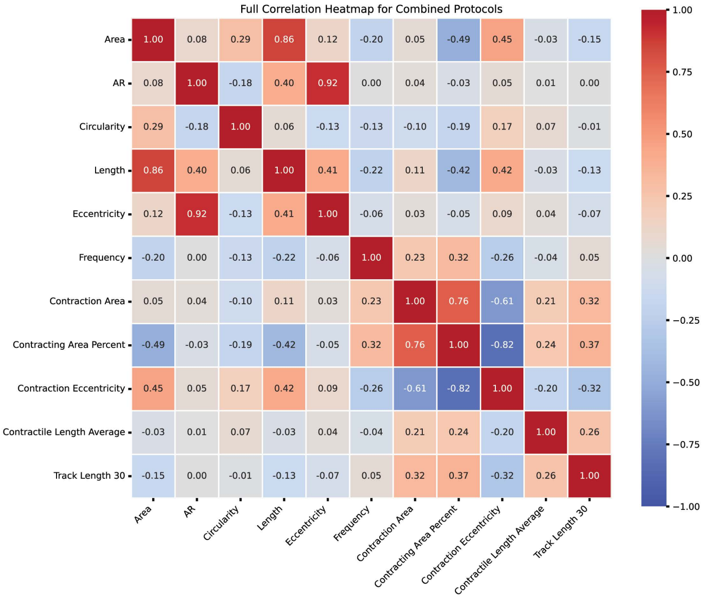


## S6B

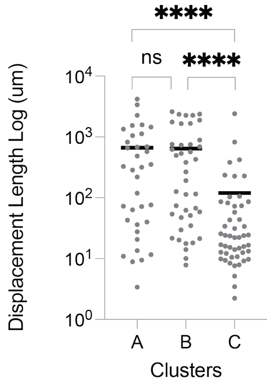


## S6C

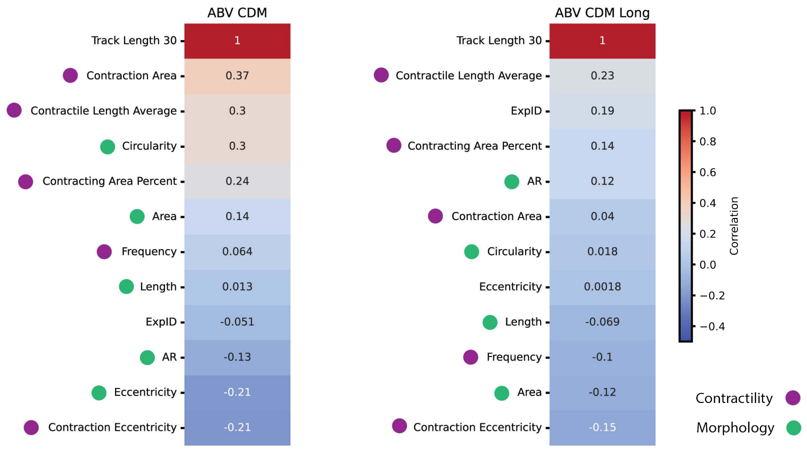

### Figure S6: Characterisation of the cluster based on their features

**A.** Correlation matrix illustrating the relationships between track length and various morphological and contractile features, including: track length, aspect ratio (AR), contraction frequency, eccentricity, average contractile length, percentage of contractile area, overall length, total area, circularity, and contraction eccentricity. Strength of correlation is reported with numbers in the matrix and with color based on the legend on the right. (*n* = 59 number of individual aggregates from cardiobot protocol, n=27 individual aggregates from cardiogenic gastruloids and n=42 individuals aggregates from extended cardiobots).

**B.** Dot plot showing displacement length (μm, $\log_{10}$ scale) for each cluster. Individual aggregates are shown, with cluster averages indicated by bold horizontal lines. Statistical analysis was performed using the Kruskal–Wallis test.

**C.** Correlation matrix illustrating the relationships between track length and various morphological and contractile features, including experiment ID for cardiobots protocol (left) and extended cardiobots protocol (right). Strength of correlation is reported with numbers in the matrix and with color based on the legend on the right. (*n* = 59 number of individual cardiobots and n=42 of individual extended cardiobots).

Statistical significance: *$p < 0.05$; **$p < 0.01$; ***$p < 0.001$
